# Neutrophil-Derived Oncostatin M Contributes to Endothelial Cell Dysfunction During *Treponema denticola* Interaction

**DOI:** 10.64898/2026.06.01.729434

**Authors:** Dayron M. Leyva-Rodriguez, Michelle B. Visser

## Abstract

Periodontitis (PD) is a common chronic inflammatory condition and a risk factor for cardiovascular diseases (CVD), yet underlying linking mechanisms remain unclear. The cytokine Oncostain M (OSM) is elevated in both PD and CVD and has emerged as a potential mediator linking oral inflammation to vascular dysfunction. Neutrophils represent a prominent source of OSM during PD and OSM production is elevated by the periodontal pathobiont *Treponema denticola* (Td). This study investigated the role of exogenous and neutrophil-derived OSM in endothelial cell (EC) dysfunction and the contribution of heterogenous oral *Treponema* species in OSM production. Human aortic endothelial cells (HAoEC) were used to evaluate the effects of exogenous purified OSM and neutrophil-derived OSM on endothelial cell function. Endothelial permeability, neutrophil transmigration, cytokine production, cell activation and junctional integrity were assessed using transwell assays, ELISAs, real-time PCR, immunoblotting and immunofluorescence microscopy. Exogenous OSM significantly increased HAoEC permeability, neutrophil transmigration and promoted endothelial activation; characterized by increased E-selectin, ICAM-1 and IL-6 expression. Mechanistically, OSM activated OSMR-STAT3 signaling and altered organization of VE-cadherin in adherens junctions and decreased expression of occludin in tight-junctions. Heterogenous oral *Treponema* species promote OSM production from mouse and human neutrophils in vitro and in vivo using a mouse air pouch model of infection. *T. denticola* most robustly induced OSM release, likely independent of prominent virulence factors dentilisin and Msp. Co-culture model experiments revealed conditioned media from *T. denticola*-stimulated neutrophils promoted endothelial cell permeability and IL-6 while reducing endothelial nitric oxide synthase (eNOS) production. These effects were abolished by antibody neutralization of OSM, supporting a casual role of neutrophil-derived OSM. Overall, these findings provide mechanistic insight into putative links between PD and adverse cardiovascular events and identify OSM signaling as critical mediator in inflammation-driven endothelial dysfunction.

## Introduction

Periodontal disease and cardiovascular diseases (CVDs) are both common conditions characterized by a chronic inflammatory state [1–5]. Periodontal disease represents a spectrum of pathologies affecting 47% of the population in the United States to some degree, while the most advanced form, periodontitis (PD), affects more than 1 billion individuals worldwide [6–8]. PD is characterized by damage and irreversible loss of tooth-supporting soft tissue and underlying alveolar bone due to an imbalance in the host immune response and a dysbiotic subgingival bacterial community. The oral cavity consists of diverse microbial community of over 700 bacterial species, and oral spirochetes including *Treponema denticola* (Td)*, Treponema maltophilum* (Tm), *and Treponema lecithinolyticum* (Tl), proliferate and thrive in this subgingival inflammatory environment during PD [9–13]. PD is unequivocally more than just an oral concern; with PD being established as a significant risk factor for a range of systemic co-morbidities, including adverse cardiovascular manifestations [14–18].

CVDs encompass a range of conditions affecting the heart and blood vessels, and are the leading cause of death worldwide, with more than 19.2 million CVD-related deaths in 2023 [19–22]. Mounting evidence has shown a strong association between CVD and PD [14–18] with periodontal therapy interventions improving outcome measures such as blood pressure, lipoprotein cholesterol levels and inflammatory markers which are known to increase future cardiovascular risk [23–25]. Despite the indications of PD influencing overall health, there remains a need to identify mediators and mechanisms contributing to CVD in the context of PD.

PD is characterized by inflammation both locally in the oral cavity and systemically throughout the body. One proposed contributor linking PD and systemic health is through a chronic inflammatory state; either by overflow of inflammatory mediators across damaged gingival tissue and/or elevated circulatory cytokine levels [4, 5, 10]. Oncostatin M (OSM) is a pleiotropic cytokine of the IL-6 family that plays a role in many biological processes and is elevated in numerous inflammatory conditions [26]. During PD, OSM is elevated locally in saliva, gingival crevicular fluid, and gingival tissue, together with increased circulating levels [2, 27–30]. While various host cells are known to produce OSM, our group has reported OSM expression and release to be increased from oral neutrophils during clinical PD [27]. The gingival tissue is under constant neutrophil immune surveillance with correlative increases in neutrophil number during PD. Neutrophils comprise 95 -98% of the leukocyte population in saliva and are the predominant innate immune cell trafficking through the gingival tissue and likely represent a primary source of OSM in the oral cavity [27, 31–33]. In murine periodontal lesions, neutrophils highly expressed OSM, which was necessary for neutrophil-osteogenic crosstalk to promote alveolar bone loss in an experimental ligature-induced periodontitis model [30]. While the oral subgingival microbial community is large, each bacterial member likely has distinct and important effects during PD. Our previous report has shown that the oral pathobiont *Treponema denticola*, distinctly promotes the secretion of OSM from neutrophils as compared to the periodontal keystone pathobiont *P. gingivalis* [27], and our recent study demonstrated that *Treponema maltophilum and Treponema lecithinolyticum* can distinctly modulate neutrophil signaling pathways to manipulate neutrophil responses [34]. We have reported OSM production in a murine air pouch model following *T. denticola* exposure [27] and *osm* gene expression is increased in the gingival tissue following polymicrobial inoculation including *T. denticola* in an experimental model of murine PD [35].

Despite the role of *T. denticola* in promoting OSM secretion from neutrophils, there remains limited knowledge of specific virulence factors responsible and whether this effect is widespread across heterogeneous oral *Treponema* species.

OSM has biphasic effects; being reported to demonstrate beneficial effects in vascular physiology, but under conditions of prolonged exposure and overexpression, OSM can have adverse effects, with worse CVD outcomes [36]. OSM is expressed in atherosclerotic lesions, is positively correlated with CVD progression, and can directly mediate changes in endothelial cells (EC) [37–40]. The endothelium composed of ECs represents the innermost layer of the vascular system, which is critical for maintaining vascular homeostasis through appropriate barrier function, vascular tone, molecular transport, immune cell adhesion, and production of cellular mediators [41, 42]. Physiological EC functions can be compromised, leading to endothelial cell dysfunction, which is well-recognized as a critical precursor event of CVDs, such as atherosclerosis [41, 43]. While endothelial dysfunction is described as changes in vascular dilation and constriction, at the endothelial cell (EC) level, it is characterized by an activated endothelium with upregulation of adhesion molecules, production of inflammatory cytokines, changes in integrity, oxidative signaling and production. Numerous risk factors, including inflammatory cytokines such as OSM, are known to drive changes in endothelial cells. For example, OSM increases gene expression of cell adhesion proteins and cytokines in ECs from diverse vascular bed origins, together with promoting endothelial activation and inflammation in an atheroprone mouse model [44]. Neutrophil-derived OSM also enhanced P-selectin clustering on EC to promote immune cell adhesion and thrombi formation in a flow-restricted vein thrombosis mouse model [45]. Despite OSM’s role in PD and CVD, no study has yet addressed how OSM may decrease vascular molecular events such as EC integrity and permeability and how neutrophil-derived OSM influences vascular EC function during *Treponema*-neutrophil interaction.

Defining initial mechanisms driving EC dysfunction in the context of OSM and during interactions with neutrophils and oral pathobionts such as *Treponema* species, are crucial for understanding pathogenic relationship between cardiovascular health and PD. Consequently, there is a crucial need to understand the role of neutrophil-derived OSM in promoting EC changes in response to *T. denticola* and other oral *Treponema* species, to define *Treponema* virulence factors associated with OSM secretion, and to understand the molecular mechanisms by which ECs may become compromised during an OSM-inflammatory environment. In this study, we used primary human aortic endothelial cells as a physiological model to represent a medium to large vessel site prone to adverse CVD events, such as atherosclerosis and thrombosis [46, 47]. In lesion prone areas, lesions begin to develop under an intact but leaky and activated endothelium layer where early pathological changes occur before detectable morphological changes in the vascular wall [48]. We hypothesized that exogenous OSM and neutrophil-derived OSM signaling during Td-neutrophil interaction disrupt endothelial junctional integrity and inflammatory signaling pathways. Our data suggest that neutrophil-derived OSM is a crucial mediator of endothelial cell dysfunction characteristics during *T. denticola* interactions, while shedding light on the mechanistic connections between periodontal disease and cardiovascular health.

## Materials and Methods

### Human Aortic Endothelial Cell Culture

Primary human aortic endothelial cells (HAoEC, PromoCell) were cultured in 12-well plates at 37 °C, 5% CO2 in endothelial cell growth medium MV (PromoCell). HAoEC were passaged using PromoCell Detachkit according to the manufacturer’s instructions. Briefly, cells were washed with Hepes (C-40010), for 15 seconds, detached using 0.04% Trypsin/ 0.03% EDTA (C-41012). Trypsin was neutralized by adding an equal amount of Trypsin Neutralizing solution (C-41110), cells were centrifuged at 220xg for 3 minutes and resuspended in fresh HAoEC medium. HAoEC between passages 3 to 7 used in this study. Once a monolayer was formed, the cells were treated with human Oncostatin M (10 ng/mL, STEMCELL CAT# 78094), human TNF-α (10 ng/mL, Fisher Scientific PHC3016,) or left untreated (control) for 24 hours as described under each individual assay section.

### Bacterial culture and cell-coculture

*Treponema denticola* (Td) type strain 35405, the dentilisin mutant strain K1, and the Msp mutant strain MHE were grown anaerobically at 37 °C in NOS media [49]. Mutant strains K1 and MHE were grown in NOS media containing 20 µg/ml erythromycin. *T. maltophilum* (Tm) type strain ATCC 51939, and *T. lecithinolyticum* (Tl) type strain ATCC 700332, were grown anaerobically at 37°C in OMIZ P4 media [49]. *Porphyromonas gingivalis* (Pg) 33277 was grown anaerobically on Blood Agar or Trypticase-Yeast Extract Broth supplemented with hemin and menadione. Spirochete cultures were examined for purity and morphology and enumerated using dark-field microscopy. For bacteria-neutrophil co-culture assays, *T. denticola* and P*. gingivalis* were grown anaerobically for 3 and 4 days, respectively, while *T. maltophilum* and *T. lecithinolyticum* were grown anaerobically for 7 days. Bacteria were then centrifuged at 2000 × g for 10 minutes at room temperature, washed once with anaerobic PBS, and resuspended in the corresponding media for the subsequent in vitro co-infection assay or exposure in murine air-pouch model.

### Western Immunoblotting

Following cell treatments, HAoEC were lysed in 100 µl PBS with 100 µl of 2X SDS sample (BioRad). All samples were boiled for 10 minutes, equal volumes (10-15µl), separated by 10% SDS-PAGE and transferred to nitrocellulose membranes. Membranes were blocked with 5% milk in TBS/0.05% Tween for 1 hour then incubated overnight at 4°C with primary antibodies **(Table 1).** Membranes were incubated with species appropriate HRP-linked secondary antibodies (1:10000) for 1 hour at RT, followed by detection with Amersham ECL Prime solutions (GE). In some experiments, HRP was inactivated with sodium azide and membranes were re-probed with the appropriate non-phosphorylated antibodies and/or anti-β-actin. Densitometry analysis was performed using ImageJ software.

**Table 1.** Primary antibodies used in this study.

| Antibody | Company | Cat. number | Dilution |
| --- | --- | --- | --- |
| OSMR | BioRad | 160411 | 1:800 |
| OSM | R&D systems | MAB295 | 100 ng/mL |
| LIFR | Santa Cruz Biotech | B0922 | 1:1000 |
| β-actin | Cell Signaling Technology | 3700S | 1:5000 |
| pSTAT3 | Cell Signaling Technology | 9145S | 1:1000 |
| STAT3 | Cell Signaling Technology | 9139S | 1:1000 |
| P-p44/42 MAPK | Cell Signaling Technology | 4370T | 1:1000 |
| p-44/42 MAPK | Cell Signaling Technology | 4695T | 1:1000 |
| pAkt | Cell Signaling Technology | 9275L | 1:2000 |
| Akt | Cell Signaling Technology | 9272S | 1:1000 |
| VE-cadherin | Cell Signaling Technology | 2500S | 1:1000 |
| β-catenin | Cell Signaling Technology | 8480S | 1:1000 |
| ZO-1 | Cell Signaling Technology | 8193T | 1:1000 |
| Occludin | Cell Signaling Technology | 91131S | 1:1000 |
| ICAM-1 | Cell Signaling Technology | 4915S | 1:500 |

### Neutrophil Isolation and Fluorescent Labeling

Human blood neutrophils from healthy donors (N=3) were isolated as described [27, 50]. Donors were recruited between January 1, 2025 to December 31, 2025, informed of the study and their consent obtained in writing under a protocol approved by the University at Buffalo Institutional Review Board (IRB Protocol Number 030-529353). This study complied with ethical standards outlined in the Belmont Report. For neutrophil isolation, peripheral blood (20 to 40 ml) was drawn from healthy donors into sodium citrate tubes (BD Vacutainer Citrate tubes). Equal volumes of blood were layered over separation gradient media (1-Step Polymorphs, Accurate Chemicals, Westbury, NY) and centrifuged according to the manufacturer’s directions. Bands corresponding to neutrophils were aspirated, washed twice with Hank’s Buffered Saline Solution without magnesium or calcium, (HBSS --, Corning, 21-022-CV) and red blood cells lysed with RBC lysis buffer (BioLegend, San Diego, CA). Neutrophils were washed with HBSS --, resuspended and counted using a hemocytometer. Neutrophil viability was assessed by trypan blue staining, and purity was ∼ 95% as assessed by morphological staining and visual observation, similar to our previous report [51]. Murine neutrophils were isolated from bone marrow of C57BL/6J mice (6 weeks old, Jackson Laboratory, Bar Harbor, ME) using Percoll density gradient separation as described [50]. Animal experiments were carried out in accordance with recommendations in the Guide for the Care and Use of Laboratory Animals under protocol approved by the UB Institutional Animal Care and Use Committee of the University at Buffalo (IACUC Protocol Number ORB07113Y). Briefly, mice were euthanized by carbon dioxide inhalation, bone marrow was flushed from femurs and tibias and fractionated on a discontinuous Percoll (Sigma) gradient (80%/65%/55%), Mature neutrophils were collected at the 80%/665% interface, washed and red blood cells lysed prior to use in assays. In some experiments, 10x10^6^ neutrophils were labeled for 20 minutes with 5 µM 5-(and -6) – Carboxyfluorescein Diacetate, Succinimidyl Ester (CFSE) (Invitrogen) according to manufacturer’s instructions and washed once with PBS without magnesium or calcium, PBS -- (21-040-CV) before use.

### Neutrophil Transmigration Assay in Pre-Treated HAoEC

HAoEC (5×10^4^) were seeded for 48 hours on a Transwell membrane (24-well plate, 3.0 μm pore size, Greiner Bio-One) followed by OSM, TNF-α, or control treatment for 24 hours. The medium containing the treatments was removed, and 3×10^5^ CFSE-labeled human neutrophils were added to the upper chamber. They were allowed to migrate for 3 hours to the receiving chamber, which contained RPMI (no fetal bovine serum (FBS) (Corning) and 1 μM N-formyl-methionyl-leucyl-phenylalanine (fMLP, Sigma) as a chemoattractant. The migrated neutrophils were collected and centrifuged at 400 xg for 10 minutes at 4°C. Neutrophils were resuspended in 50 μL of 1x HBSS (Corning), attached to coverslips (15 minutes at room temperature in the dark), and fixed by adding 50 μL of 4% PFA for 10 minutes at room temperature. Coverslips were directly mounted on glass slides with DAKO fluorescence mounting medium and fluorescent images were acquired using an Andor Dragonfly spinning disk confocal using a 63X objective attached to a sCMOS camera (Oxford Instruments Andor). Five images were collected per coverslip (middle, top-right, top-left, bottom-right, and bottom-left), and the total number of neutrophils counted using ImageJ software.

### Dextran Permeability Assays

To assess the effect of purified cytokines on endothelial cell permeability, HAoEC (1x10^5^) were grown on transwell membranes (12-well plate, 0.4 um pore size, Greiner Bio-One) for 48 hours. The apical portion of the transwell insert contained HAoEC in 500 microliters of media, and the bottom well contained 1 mL of media. After 24 hours, the apical side of the inserts were replenished with 300 uL of fresh media. Before treatment, the media from the inserts containing cells was carefully removed and inserts were transferred to a new 12-well plate with 1 mL of fresh media. Media containing OSM or TNFα (500 microliters) was added to the apical side of the well for 24 hours. Experimental test media was removed, followed by transfer of the inserts to a new 12-well plate containing 1 mL of pre-warmed phenol red-free media. FITC-dextran of 70 kDa (64 μg/mL, Invitrogen) diluted in phenol red-free media was added to the apical side of the inserts (200 μL) for 30 minutes to allow passage to the bottom receiving well at RT. Media was collected from the bottom well and 100 uL used for measurement of fluorescence intensity (485 excitation/535 emission) using FlexStation 3 Multi-Mode Microplate Reader.

To assess changes in HAOEC permeability in response to *T. denticola* -stimulated neutrophil conditioned media, PMNs (5 × 10^6^ in 1.5ml RPMI with 10% FBS) were treated with *T. denticola* (MOI 100) or sham media (control) for 3 hours in 12-well plate at 37 °C with 5% CO2. Conditioned media was collected, centrifuged (400×g for 10 minutes at 4 °C), then filtered through a 0.2 μm filter to remove bacterial remnants. Filtered conditioned media (CM) was then treated with either neutralizing OSM antibody (100 ng/mL, Table 1), goat serum as an IgG control (1:1000), or left untreated for 2 minutes at RT. Before adding CM from PMNs, media from inserts containing HAoECs was carefully removed and transferred to a new 12-well plate with 1 ml of fresh HAoEC media in the bottom well. Conditioned media treatment (diluted 1:1, 300 μl of HAoEC media and 300 μl of neutrophil C.M) was added to the top well. As additional controls, we used HAoEC media alone, 1:1 ratio of HAoEC media with RPMI +10% FBS, or 1:1 ratio of HAoEC media with CM from RPMI +10% FBS containing 5x10^8^ *T. denticola* incubated during PMN the co-infection period to assess if any unfiltered bacterial products will promote any changes in HAoEC permeability. After 24 hours of CM treatment, treatments were carefully removed, and the inserts were transferred to a new 12-well plate, and the permeability of 70 kDa FITC-dextran was assessed as above.

### Immunofluorescence of endothelial cellular junctions

HAoEC (1x10^5^) were cultured on sterilized 18 mm coverslips and allowed to form a uniform monolayer before treatment. After treatment, HAoECs were washed twice with ice-cold PBS, fixed with 4% paraformaldehyde for 10 minutes at room temperature (RT), permeabilized with 0.1% Triton X-100 for 5 minutes at RT followed by blocking with 0.5% Bovine serum albumin (BSA) for 30 minutes. Primary antibodies directed against the adherens junction components were incubated with cells for 2 hours at room temperature (RT), followed by a secondary antibody and actin staining with phalloidin (Invitrogen) and nuclei staining with DAPI (Roche) (Table 1). Fluorescent images were acquired using an Andor Dragonfly spinning disk confocal using a 63X objective attached to a sCMOS camera (Oxford Instruments Andor). At least five images were taken per coverslip using the same setting (Exposure time and Laser Intensity) across all samples.

### Quantitative RT-PCR

Following treatment, cells were washed with cold PBS, and RNA was isolated using E.Z.N.A Total RNA Kit I (OMEGA). RNA (100ng) was converted to cDNA using the iSCRIPT kit (Biorad), and qPCR was performed with 1μl of cDNA template in a 10 μL reaction using the SSOSyber kit (Biorad) on an Applied Biosystems 7500 machine or CFX OPUS96 Real Time PCR Machine, Bio-Rad. Relative fold change (ΔΔCt) was determined by normalizing to the housekeeping gene GAPDH in the untreated samples (control) using the primer set as previously reported [52]. Primer sequences were chosen based on literature or designed at the OriGene website as listed in Table 2 and synthesized by Eurofins Genomics (Louisville, KY).

**Table 2.**
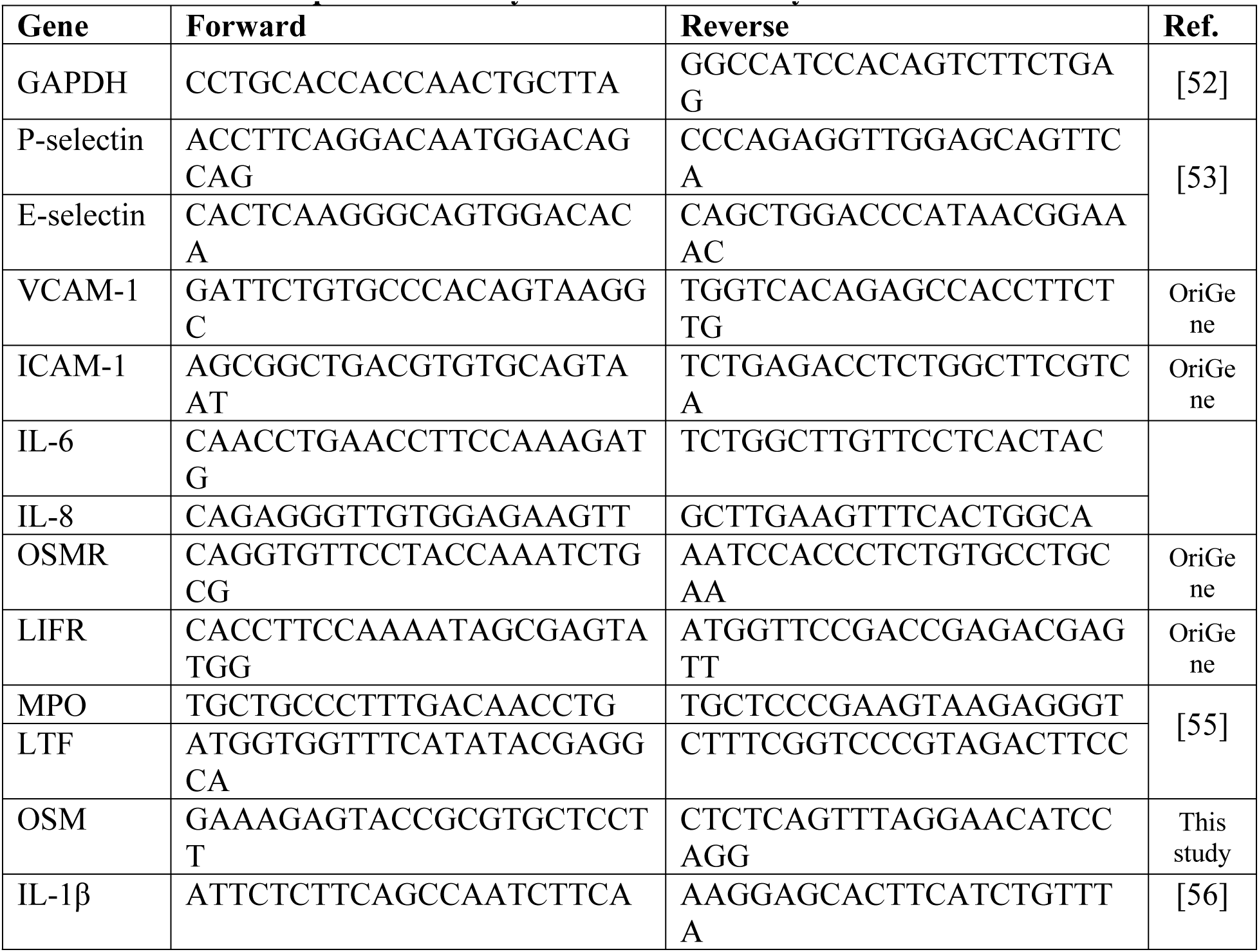
Primers for qRT-PCR analysis used in this study.

### Air pouch model of infection

A murine air pouch model of inflammation was performed as previously described [27, 57] as approved by the University at Buffalo’s Institutional Animal Care and Use Committee (IACUC ORB07113Y) and carried out in accordance with recommendations in the Guide for the Care and Use of Laboratory Animals. In brief, air pouches were formed on the dorsal region of 6-week-old C57BL/6J male mice (n=4 mice per group, Jackson Laboratories, Bar Harbor, ME) anesthetized using isoflurane. Air pouches were formed by subcutaneous injection of 3 mL of sterilized air on Day 1 with reinflation with 2 mL of sterilized air on Day 3. On Day 5, 1 × 10^9^ bacteria in 1 mL of PBS were injected into the air pouch, whereas sham mice were injected with 1 mL of PBS alone. Six hours postinfection, mice were euthanized by CO_2_ inhalation, and air pouches were washed with 2 mL of PBS. Lavage fluid was centrifuged (2000 × g for 5 min) to remove cells and then further centrifuged at 13,000 × g for 1 minute at 4°C to remove cellular debris and stored at -80 °C for ELISA analysis.

### Cytokine measurement by ELISA

Cytokines produced in lavage fluid or conditioned cell culture media were measured using commercial ELISA kits. For in vitro experiments, 5×10^6^ neutrophils (human or murine) were co-incubated with bacteria (multiplicity of infection, MOI =100) in 3 mL final volume of RPMI + 10% FBS media for 3 hours in 6-well plates at 37 °C with 5% CO2 before collecting the conditioned media. GM-CSF (100ng/mL) was used as a positive control. Conditioned media containing cellular products was centrifuged at 5000 × g for 2 minutes at 4 °C to remove cellular debris and then frozen at -80 °C until use. For OSM measurement, conditioned media from neutrophil-bacterial co-infection were concentrated using Amicon Ultra-0.5 mL centrifugal filters (molecular weight cutoff of 3000Da). Collected murine airpouch lavage fluid samples were measured directly. Cytokines (mOSM (R&D DY495-05), hOSM (R&D DY295), hIL-8 (R&D DY208), and hIL-6 (R&D DY206)) were measured by ELISA using 100 μL of media per well in duplicate, following the manufacturer’s instructions. Results were calculated using a standard curve and quantified as changes in densitometry, as measured with a FlexStation 3 Multi-Mode Microplate Reader.

### Statistical Analysis

Comparisons between two groups were performed using paired or unpaired t-tests, as appropriate. Comparisons between more than two groups were performed using ANOVA with post hoc Tukey HSD or Dunn’s multiple comparison tests, as appropriate. All statistical analyses were performed using GraphPad PRISM software (GraphPad). In vitro results are based on at least three independent experiments, whereas in vivo experiments consisted of three to four animals per experimental group. Statistical significance was defined as a P value of less than 0.05. Error bars represent the standard error of the mean (SEM).

## Results

### Exogenous OSM increases HAoEC permeability and neutrophil transmigration

EC dysfunction plays a crucial role in the development of CVDs, with increased barrier permeability characteristic of dysfunctional EC monolayers [41]. As circulating levels of OSM are elevated in both periodontal diseases and CVDs [28, 29, 58, 59], we sought to determine whether exogenous OSM exposure to HAoEC would promote changes in permeability using a FITC-Dextran permeability assay. Dextran permeability assay demonstrated significantly increased permeability across HAoEC monolayers after 24 hours of treatment with OSM or TNF-α, as evidenced by increased fluorescence intensity (Fig 1A). Along with changes in barrier permeability, increased adhesion and migration of leukocytes are characteristics of initial EC dysfunction and critical features leading to an atheroprone environment [60, 61]. A neutrophil migration assay was performed to determine if OSM would also facilitate leukocyte migration. OSM- or TNF-α-treated HAoEC demonstrated higher neutrophil migration across the monolayer as compared to sham media control exposure (Fig 1B and C). Together, these data indicate that OSM exposure promotes changes in HAoEC characteristic of initial EC dysfunction.

**Fig 1.**
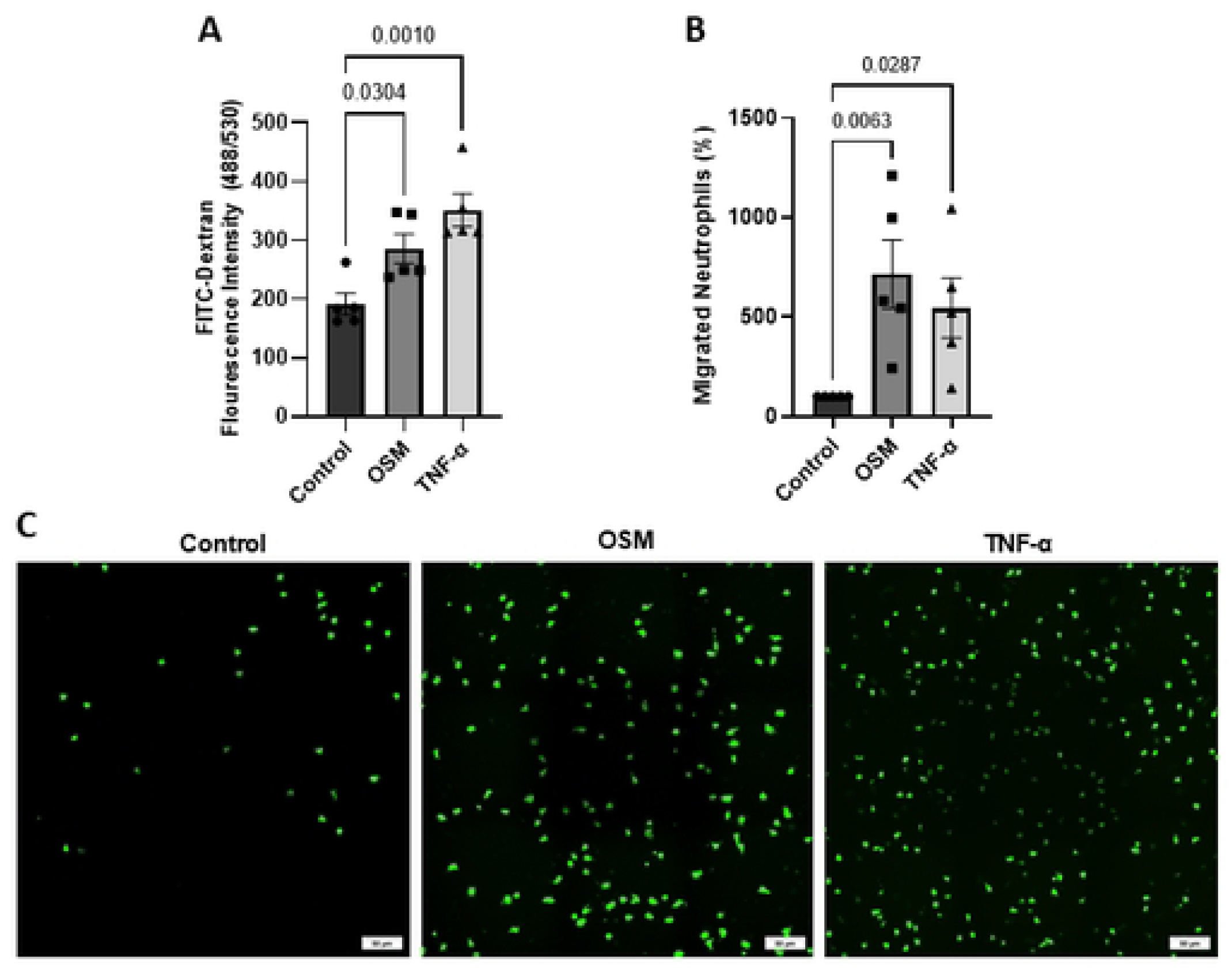
Exogenous OSM increases permeability and neutrophil migration across HAoEC. HAoEC monolayers were treated with hOSM or hTNF-a (10ng/mL), or sham media (control) for 24h on a transwell insert followed by the addition of FITC-dextran to the upper compartment for 30 minutes or CFSE-labeled human neutrophils (green) for 3h. **A**) FITC-dextran fluorescence measurements in the bottom transwell. **B**) Quantification of migrated neutrophils across HAoEC monolayers in transwell towards fmlp as a chemoattractant. Representative images of transmigration of human neutrophils (green) across HAoEC monolayers are shown in panel **C**. Bar = 50 μm. Graphs represent the mean ± SEM, n ≥ 3, p values are shown.

### Exogenous OSM supports an active inflammatory endothelial environment

Given the observed increase in permeability and neutrophil transmigration across HAoEC following OSM treatment, we next sought to assess alterations in key molecular processes underlying neutrophil transmigration. We first examined the gene expression of endothelial adhesion molecules (selectins and Cell Adhesion Molecules (CAMs)), which are responsible for initial rolling and adhesion steps of leukocyte transmigration and are indicative of an activated endothelium [41]. Gene expression of E-selectin and intercellular adhesion molecule (ICAM-1), but not P-selectin or vascular cell adhesion molecule (VCAM-1), were significantly increased in OSM-treated HAoEC compared to the control (Fig 2A). An inflammatory response is characterized by two major events: leukocyte recruitment to the affected tissue and increased vascular wall permeability, both of which can be mediated by ICAM-1 on ECs [62]. Since ICAM-I is upregulated by inflammatory mediators such as TNF-α, and given its known role as a modulator in CVDs [63, 64], we also examined its expression at the protein level. Western immunoblot analysis revealed a significant increase in total ICAM-1 protein levels in HAoEC following OSM exposure when compared to control (Fig 2B). We then assessed the effect of OSM on select pro-inflammatory mediators in HAoEC by examining both IL-6 and IL-8 gene expression and protein production levels. OSM-treated HAoEC significantly increased both IL-6 gene expression and secretion compared to the control group (Fig 2C and D), while IL-8 expression and secretion were decreased considerably (Fig 2E and F). Overall, these data support OSM in promoting cellular and molecular changes, as well as cytokine changes conducive to a pathogenic endothelial environment.

**Fig 2.**
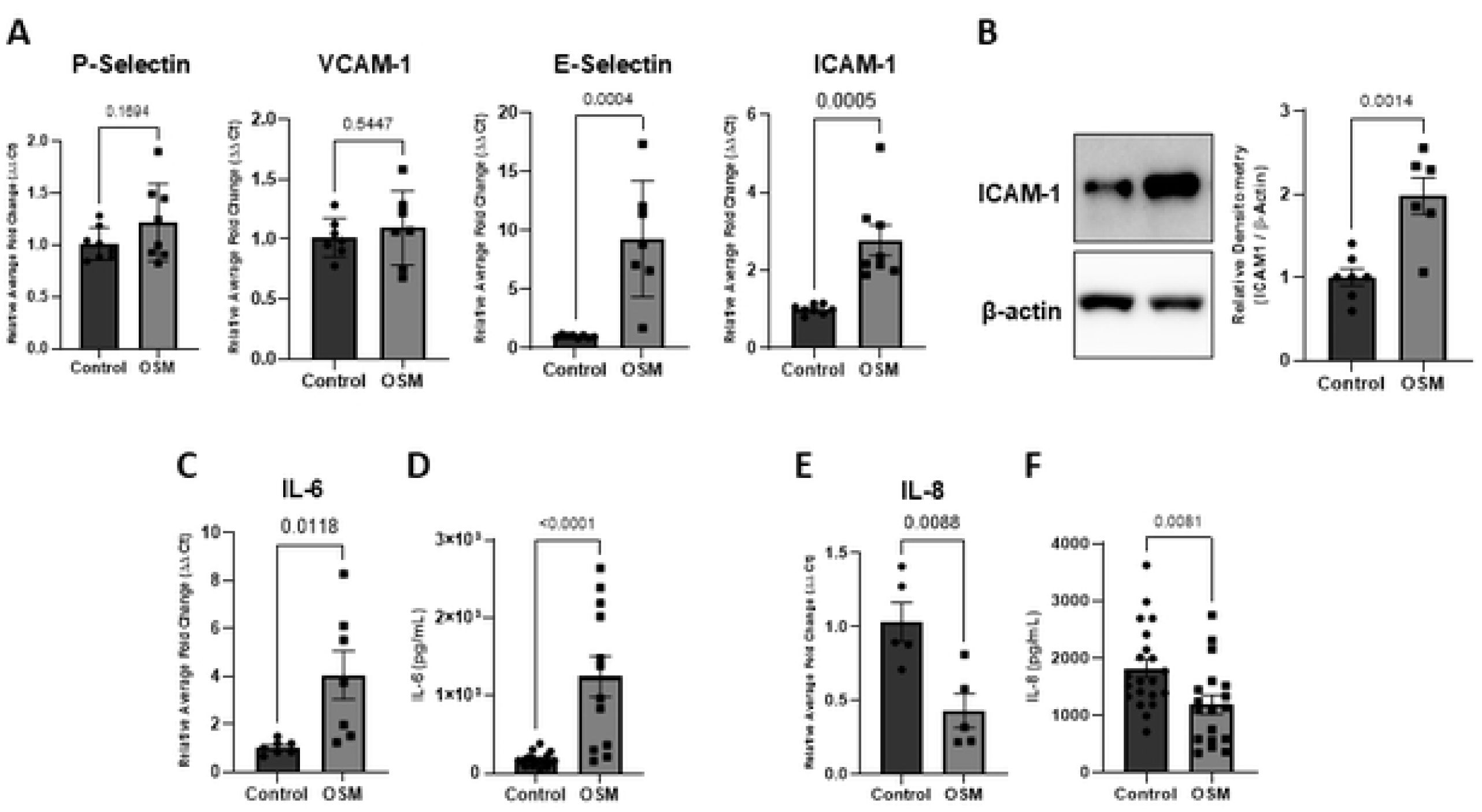
OSM can promote an increase in inflammatory mediators and activation markers in HAoEC. HAoEC were treated with OSM or no treatment (Control) for 24h. **A**) Gene expression of adhesion molecules in HAoEC (RT-PCR). **B**) ICAM-1 protein levels in HAoEC. Representative immunoblot and relative densitometry of multiple experiments are shown. Gene expression (RT-PCR), **C** and **E**, and secreted levels (ELISA) of **D**) IL-6 and **F**) IL-8. All graphs represent the mean ± SEM, n ≥ 3, p values are shown.

### OSM supports OSMR-pSTAT3 signaling in HAoEC

In humans, OSM is a ligand for two distinct receptors: the Oncostatin M Receptor (OSMR) and Leukemia Inhibitory Factor Receptor (LIFR) [65]. As both receptors have been reported to contribute to different forms of CVDs [66, 67], we next investigated which pathway is engaged in HAoEC following OSM exposure. After 24 hours of exogenous OSM treatment, OSMR gene expression was significantly increased as compared to the control (Fig 3A), but no consistently significant change in total protein level was observed (Fig 3B). On the other hand, LIFR gene expression was unaffected by OSM treatment (Fig 3C), yet a significant reduction in total LIFR protein levels was demonstrated (Fig 3D). To investigate signaling pathways downstream of OSM receptor engagement in HAoEC, we selected key components of primary pathways known to be regulated by OSM. Western immunoblot analysis revealed that phosphorylation of STAT3, but not MAPK or AKT, is significantly increased (Fig 3E, F, and G), suggesting that OSM primarily activates OSRMR-STAT3 signaling in HAoEC.

**Fig. 3.**
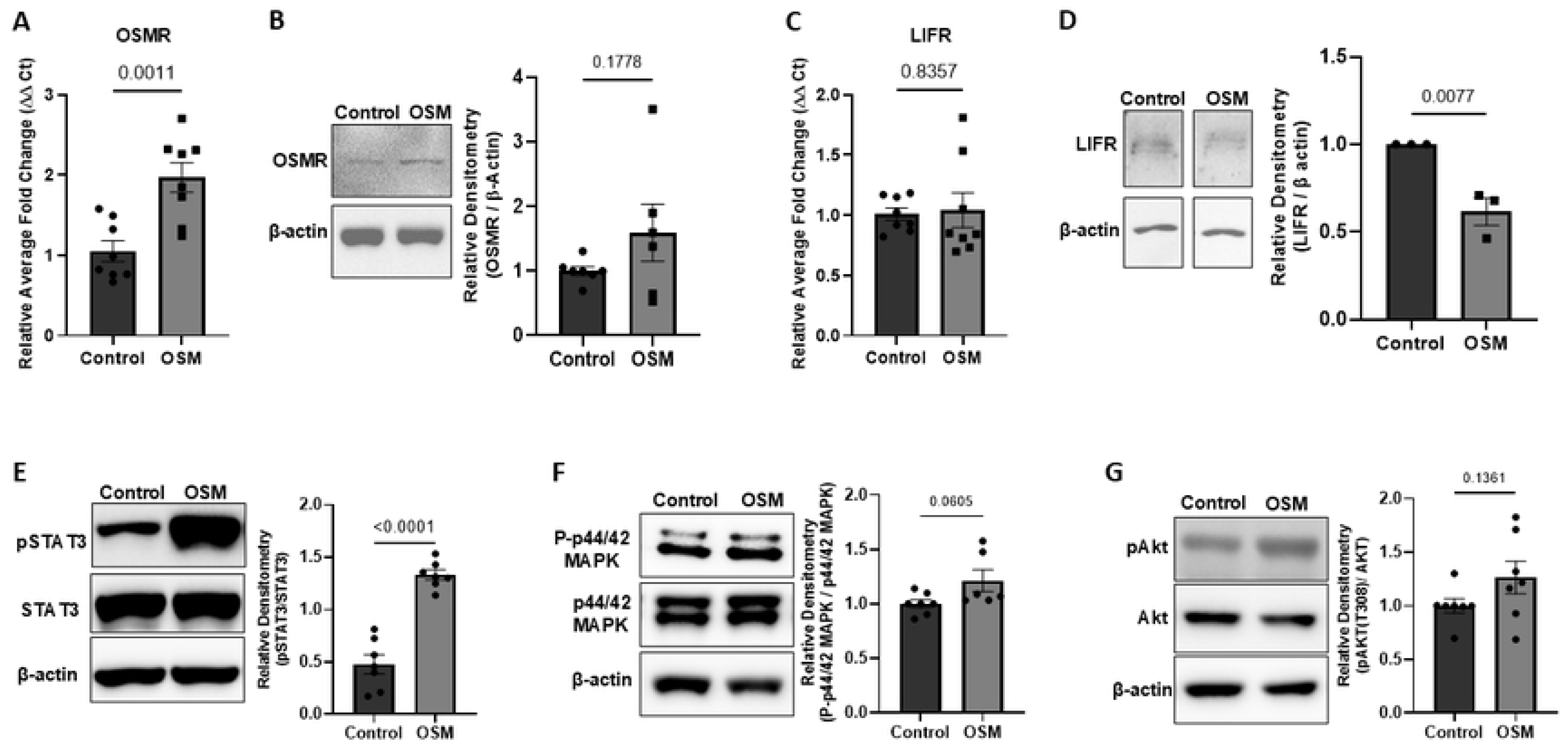
OSM promotes OSMR expression and pSTAT3 activation in HAoEC. HAoEC were treated with OSM or no treatment (Control) for 24h. Gene expression, representative immunoblots, and relative densitometry of **A and B**) OSMR and **C and D**) LIFR. Representative immunoblots and relative densitometry of **E**) pSTAT3, **F**) p44/42MAPK, **G)** pAkt. Graphs represent the mean ± SEM, n ≥ 3, p values are shown.

### OSM may contribute to a compromised HAoEC monolayer via molecular changes of VE-cadherin and occludin

An increase in endothelial permeability and leukocyte transmigration can be attributed to molecular changes in cell-to-cell adherens junction and tight junction complexes [41, 68]. To evaluate these complexes during exogenous OSM exposure, we performed western immunoblots to assess total protein expression of representative molecules. OSM-treated HAoEC showed decreased levels of the tight junction transmembrane protein occludin, yet no differences in total levels of the cytoplasmic linker ZO-1 (Fig 4A and B). OSM exposure did not significantly change the total protein levels of the adherens junction transmembrane protein VE-cadherin or the cytoplasmic linker β-catenin (Fig 4C, D). Stability of adherens junctions is provided by VE-cadherin and alterations in protein localization and interacting intracellular partners also contribute to changes in vascular permeability [69, 70]. Therefore, we performed immunofluorescence (IF) to assess morphological changes in VE-cadherin during OSM exposure. Interestingly, OSM-exposed HAoEC showed significantly increased overall fluorescence intensity and increased intracellular extensions of VE-cadherin staining when compared to control cells, which show a more linear intercellular pattern of staining (Fig 4E). These data suggest that a compromised HAoEC monolayer following OSM exposure is associated with changes in tight junction protein occludin and alterations in VE-cadherin distribution, supporting increased leukocyte migration and permeability in HAoEC during an OSM inflammatory environment.

**Fig 4.**
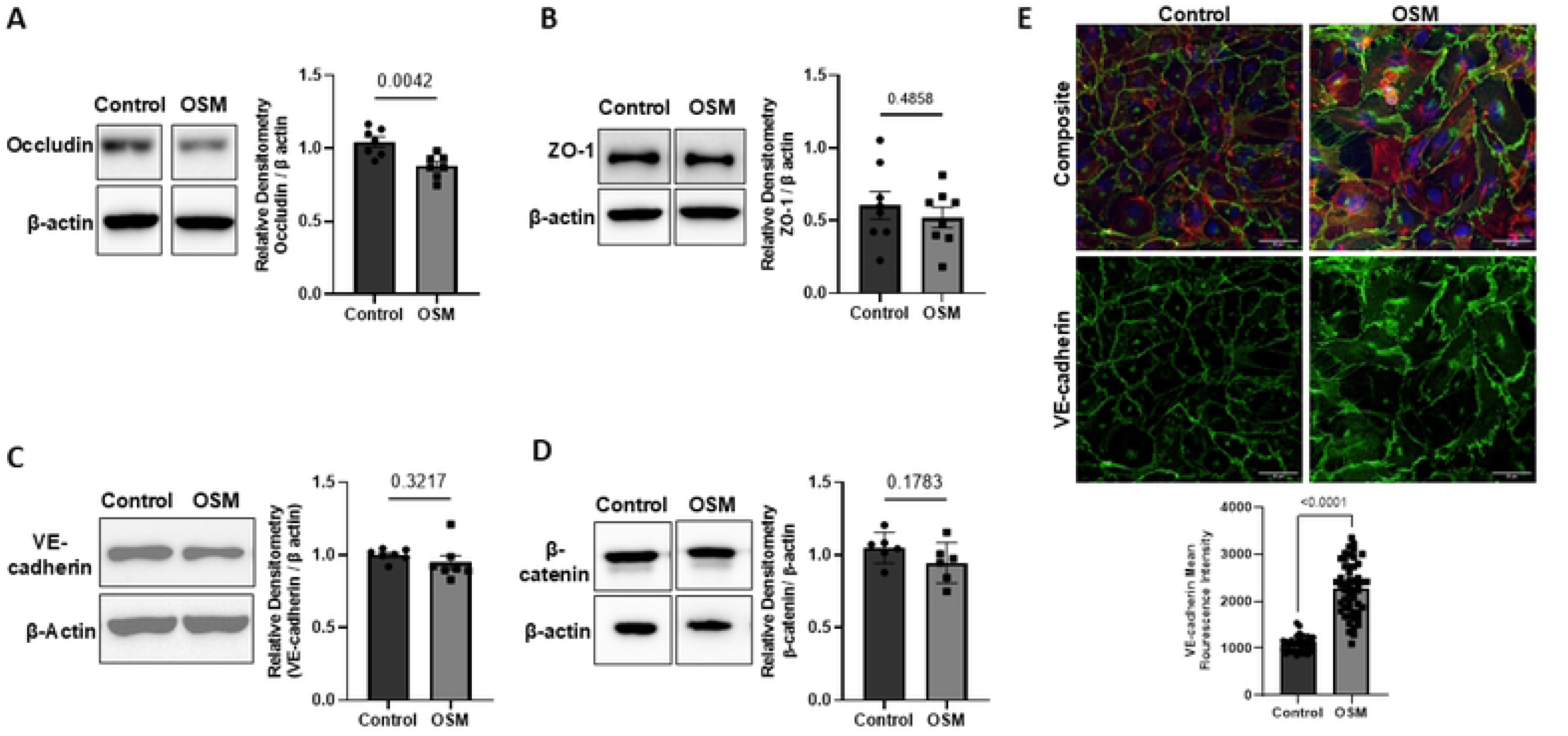
OSM contributes to the molecular changes of adherens VE-cadherin and tight junction protein Occludin in HAoEC. HAoEC were treated with OSM or no treatment (Control) for 24h. Representative immunoblots and relative densitometry of tight junction proteins **A**) Occludin, **B)** ZO-1 and adherens proteins **C**) VE-cadherin, **D**) β-catenin. **E**) Immunofluorescence images of VE-cadherin (green), actin (red) and DAPI (blue) staining in HAoEC. Graph represents the VE-cadherin mean fluorescence intensity. Bar = 50 μm. All graphs represent the mean ± SEM, n ≥ 3, p values shown.

### *T. denticola* is the primary *Treponema* species promoting OSM secretion

Our previous work has demonstrated that *T. denticola* promotes OSM release from preformed granules and de novo synthesis in PMNs [27], yet specific bacterial components triggering this remain unknown. Hence, we assessed whether prominent *T. denticola* virulence factors contributed to OSM production in PMNs. We measured released OSM levels in conditioned media via ELISA after PMNs exposure to wildtype *T. denticola*, the dentilisin protease-mutant strain K1 (lacking prolyl-phenylalanine specific protease activity), or the Msp-mutant strain MHE (lacking the major outer sheath protein, Msp) for 3 hours. Interestingly, PMN-derived OSM levels following exposure to either mutant strain was not significantly different from those following wild-type exposure, suggesting that OSM secretion from PMNs is independent of Msp and dentilisin (Fig 5A).

**Fig 5.**
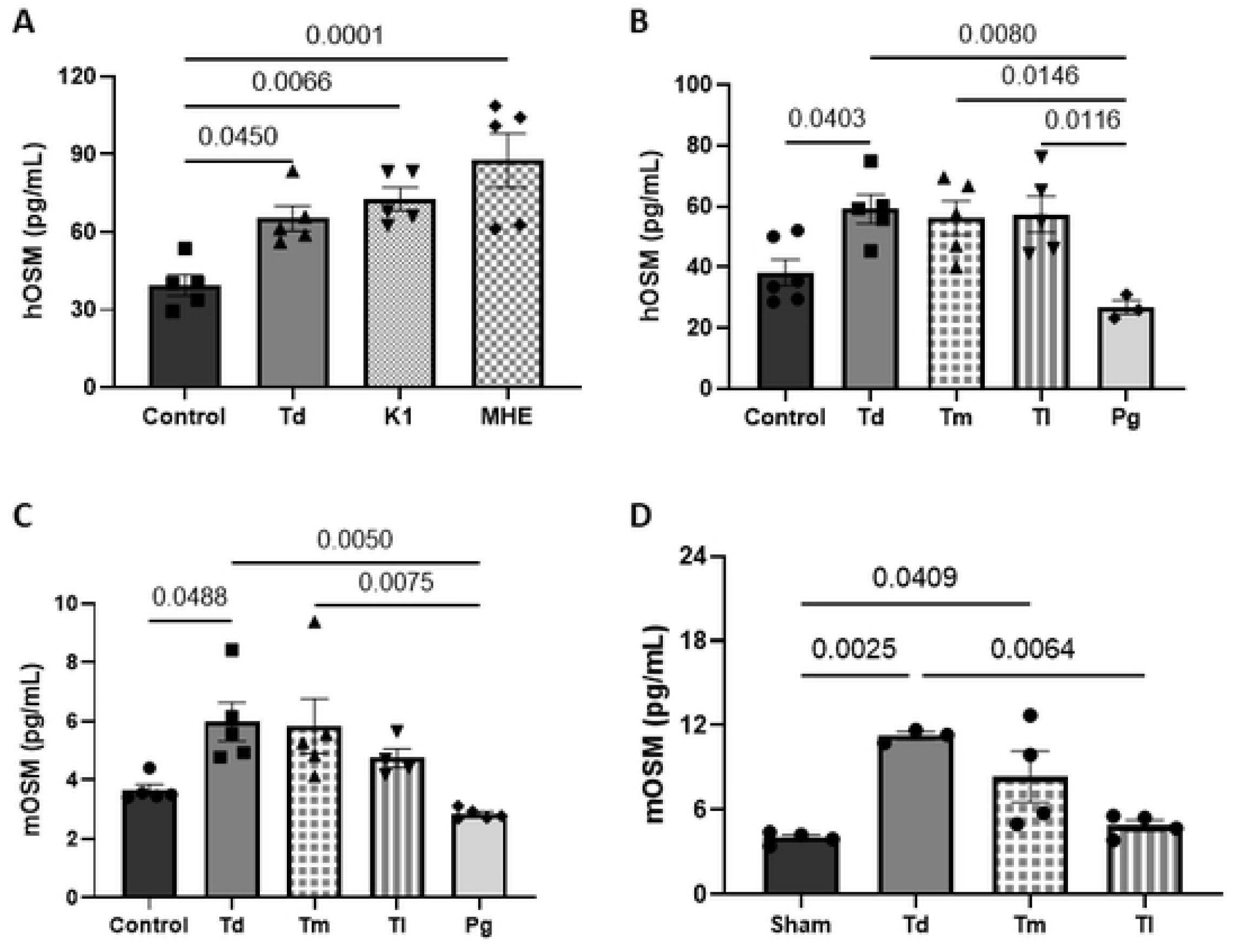
*Treponema species* promote neutrophil OSM levels in vitro and in an air pouch model of exposure. Human neutrophils were co-incubated for 3 hours at a multiplicity of infection (MOI) of 100 with various treatments: **A**) no treatment (control), wild-type bacteria *Treponema denticola* (Td), mutant bacteria K1 (which lacks prolyl-phenylalanine-specific protease activity), or MHE (which lacks the major outer sheath protein, Msp); or **B**) Td, *Treponema maltophilum* (Tm), *Treponema lecithinolyticum* (Tl), or *Porphyromonas gingivalis* (Pg). OSM levels in conditioned media were measured by ELISA. **C**) Murine neutrophils were co-incubated for 3h,3 hours (MOI 100) with Td, Tm, Tl, Pg, or no treatment (control), and OSM secretion was measured via ELISA. **D**) Air pouch lavage fluid of OSM levels were measured 6 hours after murine exposure to Td, Tm, Tl, or no treatment (sham) by ELISA. Each data point represents an independent mouse within each treatment group. All graphs represent the mean ± SEM, with n ≥ 3, p values shown.

During periodontal disease progression, heterogenous *Treponema* species become elevated in the oral biofilm community [71, 72], including *T. denticola* and the understudied species *T. maltophilum* and *T. lecthinolyticum*, which we have recently shown can modulate neutrophil function and signaling through distinct species-specific mechanistic processes [57]. Human and murine neutrophils are known to have species-specific differences in their proteome and transcriptomic landscapes [73, 74] which can represent distinct functionality. Together, this information led us to investigate whether OSM secretion from neutrophils is a unique characteristic of *T. denticola* and whether human and murine neutrophils exhibit similar responses. To interogate these questions, human or murine PMNs were exposed to *T. denticola* (Td), *T. maltophilum* (Tm), *or T. lecithinolyticum* (Tl), and *P. gingivalis* (Pg) for 3 hours, followed by measurement of OSM in conditioned media. While all three *Treponema* species elevated OSM release from both human and murine neutrophils in vitro, only *T. denticola* promoted a significant increase of OSM release compared to control neutrophils (Fig 5B and C). Similar to our previous report [27], *P. gingivalis* did not significantly change OSM production in human or murine neutrophils compared to control exposure (Fig 5B and C).

To further investigate the ability of heterogenous oral treponemes to promote the secretion of OSM using a physiological *in vivo* biological system, we employed an air pouch model of infection. Following 6 hours of exposure, increased OSM levels in lavage fluid were observed for *T. denticola* and *T. maltophilum* compared to the sham group (Fig 5D). In contrast, *T. lecithinolyticum* did not promote a significant increase in OSM release in the air pouch (Fig 5D). Overall, this data shows that heterogenous *Treponema* species can promote OSM release from both murine and human neutrophils in vitro and in an in vivo physiological model; with *T. denticola* more robustly promoting OSM release during direct neutrophil interactions and in a multicellular complex biological environment.

### Neutrophil-derived OSM is a major contributor in promoting EC dysfunction during *T. denticola*-PMN interaction

As our data herein indicates that exogenous OSM increases the permeability of human aortic endothelial cells, and our previous work demonstrated that *T. denticola* promotes OSM production and synthesis from neutrophils [27], we wanted to assess the contribution of PMN-derived OSM in promoting EC dysfunction features in the context of Td stimulation. We utilized a co-culture model together with an OSM-specific neutralizing antibody where neutrophil conditioned media (CM) from *T. denticola* or sham exposure was collected and incubated with HAoEC followed by selected downstream cellular assays. Similar to our previous report, [27], we observed a significant increase in OSM levels in the CM of neutrophils exposed to *T. denticola* compared to sham treatment (Fig 6A). EC dysfunctional changes include altered barrier permeability, and we have shown that exposure to exogenous OSM increases HAoEC permeability of FITC-Dextran (Fig 1A). Exposure of HAoEC to CM from PMNs exposed to Td (Td-PMN-CM) significantly increased passage of dextran across the monolayer as compared to exposure to control neutrophil CM (PMN-CM) (Fig 6B), indicating that neutrophil products released in response to *T. denticola* interaction promote changes in endothelial permeability. To specifically interrogate the effect of neutrophil-derived OSM in this system, Td-PMN-CM was treated with OSM-neutralizing antibody or serum (as an IgG control) prior to exposure to HAoEC. In contrast to Td-PMN-CM or IgG-treated CM (Serum-Td-PMN-CM), Td-PMN-CM exposed to an OSM neutralizing antibody (anti-OSM Td-PMN-CM) did not show a significant increase in permeability (Fig 6B). To confirm that these changes are exclusive to PMN secreted products, exposure to culture media containing only *T. denticola* during the co-culture period did not change permeability compared to control treatment. Given the notable levels of OSM secretion we observed in neutrophil CM (Fig 6A), we hypothesized that differing OSM levels would correlate with increased permeability in HAoEC. In fact, our preliminary correlation analysis with the small number of samples in this in vitro experiment revealed a strong positive correlation between increasing OSM levels and permeability (Fig 6C).

**Fig 6.**
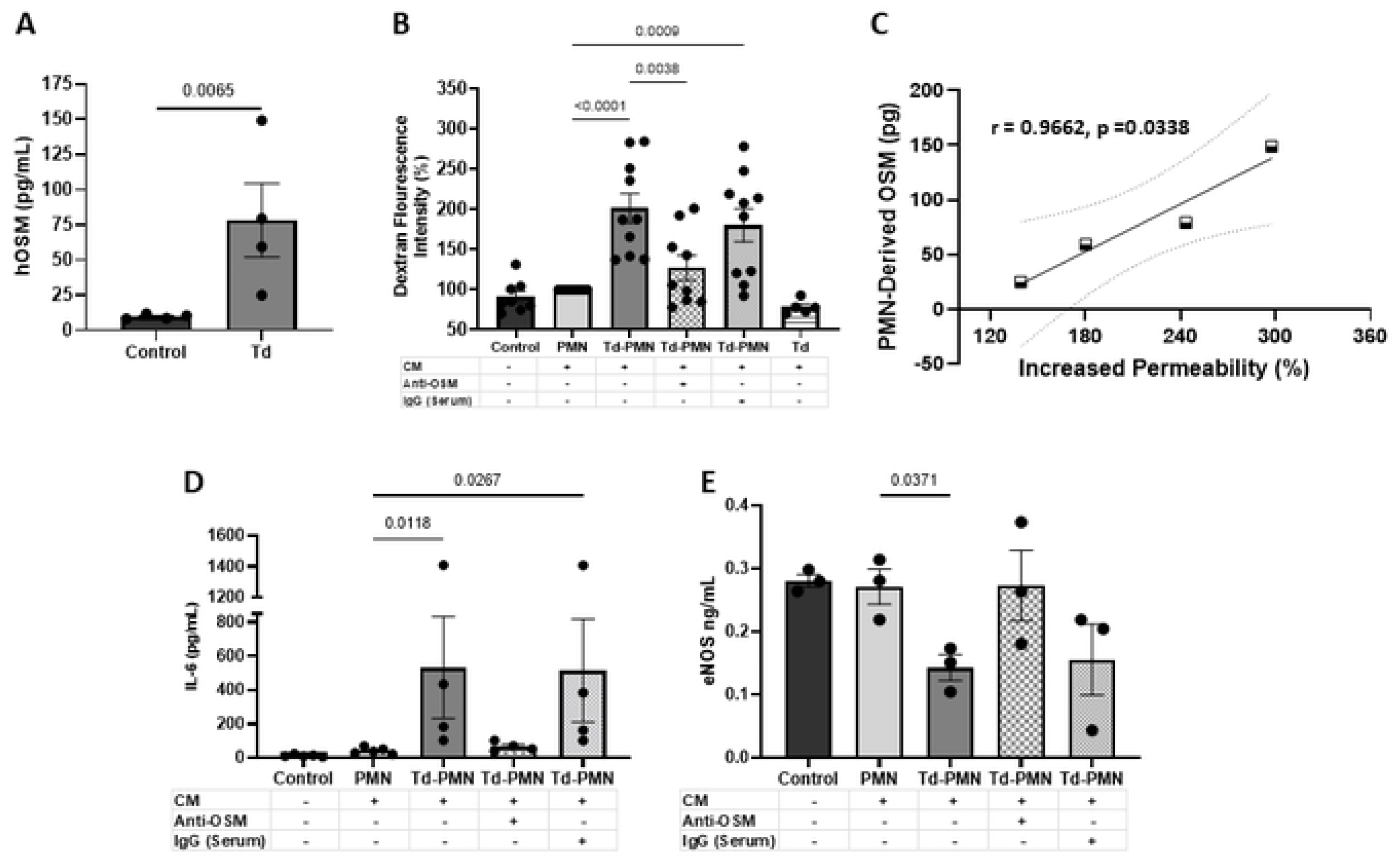
Neutrophil-derived OSM contributes to altered endothelial cell permeability and inflammatory profiles during *T. denticola*-PMN interaction. PMNs were co-incubated with *T. denticola* (MOI 100) or control (no treatment) for 3 hours. Cell-free conditioned neutrophil supernatants were collected and exposed to EC with or without OSM-neutralizing antibody (anti-OSM), or serum (IgG control). **A**) OSM secretion levels from PMNs measured by ELISA. **B**) Permeability assay assessed by FITC-dextran passage across HAoEC monolayers after 24 hours of treatment with Pearson correlation analysis of OSM levels and permeability relationship (r=0.9662, p = 0.0338) shown in panel **C**. **D**) IL-6 and **E)** eNOS production levels in HAoEC after 24 hours of treatment with PMN-conditioned media. Graphs represent the mean ± SEM, with n ≥ 3, p values shown.

We next asked how *T. denticola*-stimulated neutrophil-derived OSM contributes to production of secreted mediators characteristic of a dysfunctional endothelium. In this study, we demonstrated that exogenous OSM elevated IL-6 production from HAoEC (Fig 2B) and thus selected this as a representative cytokine for measure. Like exposure to exogenous OSM, exposure to Td-PMN-CM or Serum-Td-PMN-CM significantly increased IL-6 protein levels from HAoEC as compared to PMN-CM alone (Fig 6D). However, treatment of Td-PMN-CM with OSM-neutralizing antibody prior to exposure with HAoEC reduced IL-6 production to levels comparable to those in cells treated with PMN-CM, indicating the role of PMN-derived OSM in promoting an inflammatory environment.

Nitric oxide (NO) is an essential molecule produced by endothelial cells via endothelial nitric oxide synthase (eNOS), which plays a crucial role in cardiovascular health [75–77]. Reduced eNOS activity is directly linked to compromised endothelial monolayers and CVDs [78–80], thus we wanted to investigate the impact of PMN-derived OSM on this pathway. Secreted eNOS levels from HAoEC were significantly reduced by exposure to Td-PMN-CM as compared to PMN-CM, which was reversed by treatment with OSM neutralizing antibody (Fig 6E). Together, these data confirm a role for PMN-derived OSM in facilitating an environment conducive to CVD development after Td interaction; by both modifying the HAoEC inflammatory environment through increased IL-6 secretion, and reduced eNOS production and functionally by promoting EC permeability.

## Discussion

Oncostatin M has emerged as an essential mediator associated with both CVDs and PD [1, 28, 30, 32, 37–40, 44, 81]. In this study, we extend knowledge that exogenous OSM directly promotes pathogenic endothelial features including neutrophil transmigration and loss of junctional integrity. We also demonstrate that neutrophil-derived OSM is a key mediator linking *T. denticola* exposure to pathological endothelial barrier changes. These findings provide mechanistic insight into how oral organisms may contribute to both local and systemic vascular pathology.

Vascular inflammation and endothelial cell activation represent initial pathogenic changes occurring before any detectable morphological changes in the vessel walls. In this present study, exogenous OSM or neutrophil-derived OSM produced during *T. denticola* interaction induces aortic endothelial cell activation and proinflammatory cytokine signaling. In line with previous reports [44, 81], OSM induced upregulation of endothelial cell adhesion molecules E-selectin and ICAM-1. E-selectin is found exclusively in endothelial cells, significantly increased in inflammatory environments and regulates attachment and stable adhesion of leukocytes to the endothelium in various disorders, such as CVDs [82–84]. A non-significant trend of increased P-selectin gene expression was observed in our study. While initially this seemed unexpected as P-selectin is crucial for initial neutrophil adhesion and endothelial cell activation, this may indicate *in vitro* dose, time-dependent or endothelial cell origin dependent responses. OSM has been reported to promote P-selectin gene and protein expression in HUVECs and HDMECs and P-selectin clustering in HUVECs [44, 45, 85]. We also did not observe significant changes in VCAM-1 gene expression following OSM exposure, but this could be due to difference in analysis points selected as this appears time-dependent in HAoECs. The observed increase in ICAM-1 at both the gene and protein levels is notable, given its established role in leukocyte arrest and association progression of adverse cardiovascular events [62, 63, 86–88]. Our in vitro findings are in line with reports that ICAM-1 protein expression is elevated in the aortic root of hyperlipidemic mice following chronic 3-week OSM exposure [44].

Concurrently, OSM exposure increased HAoEC IL-6 production while suppressing IL-8 expression and secretion. Elevated circulating levels of IL-8 have been associated with adverse cardiovascular outcomes together with increased expression in vessels of diverse origin and vascular disease conditions. Our data indicating that OSM decreased both IL-8 gene and protein production from HAoEC in vitro suggests this cell type may not be a primary source of IL-8. In line with our findings, OSM upregulated cytokine expression including IL-6 in human cerebral endothelial cells and HUVECs yet IL-8 remained unchanged [81, 89]. Synergistic or antagonistic effects between OSM and other cytokines or inflammatory products has been reported in other cellular systems. For example, OSM and LPS synergize to increase MCP-1, IL-6 and VEGF yet IL-8 levels are reduced in human aortic adventitial fibroblasts and smooth muscle cells [90],while OSM and TNFα combined to enhance IL-6 and MCP-1 production but downregulate IL-8 expression [89]. Mechanistically, this could involve selective receptor engagement or transcription factor-promotor interaction and needs to be investigated in HAoECs as well as in a more complex multicellular system.

IL-8 (CXCL8) is well established to be a classical leukocyte chemokine, yet our data indicates OSM induces significant neutrophil transmigration across HoAEC monolayers without IL-8 expression or release. It has been reported that OSM can also induce synthesis of other neutrophil chemokines such as CXCL1 (GROα), CXCL2 (GROβ) and CXCL5 (ENA-78) from HUVECs [81], thus these could be significant contributors to OSM-mediated neutrophil interaction in this system. While elevated IL-8 levels have been reported in saliva or gingival crevicular fluid (GCF) of subjects with periodontitis [91, 92], contradictory studies have also reported that IL-8 levels may be decreased locally in the gingival crevicular fluid (GCF) [92–94] as well as in serum during periodontitis [95]. This conflicting data could suggest why some individuals may be more susceptible to periodontal disease if IL-8 neutrophil recruitment and activity is ineffective. We and others have reported OSM is elevated during PD [2, 28] and it would be interesting to investigate potential associations between circulating OSM and IL-8 levels with both oral health and cardiovascular status.

EC activation can be driven by STAT3 signaling via OSMR and LIFR during interaction with OSM [44]. Our findings indicate that OSM preferentially engages the OSMR–STAT3 axis while decreasing LIFR protein levels in HAoEC. STAT3 signaling has been implicated in various cardiovascular conditions such as atherosclerosis [96] and aortic aneurysm [97]; and modulating STAT3 signaling has potential as a valid therapeutic option for modulating initial pathogenic events of vascular disease [98]. Forthcoming work should examine the spectrum of STAT3-regulated genes in HAoEC, as well as exploring how co-stimulation with other inflammatory cytokines influences receptor dynamics and downstream outcomes, to increase the understanding of OSM’s role in vascular pathology. The LIF-LIFR axis may still play a role in vascular disease as single-cell analysis of human carotid plaques revealed increased LIF expression together with LIFR expression in activated EC [66]. Dysregulation of LIFR signaling has been implicated in various diseases and LIF can play a pro- or anti-inflammatory role dependent on tissue context and microenvironment. Mouse model studies have revealed that small-molecule inhibition of LIFR reduced atherosclerosis outcomes while overexpression suggests that LIF signaling can have cardioprotective effects in myocardial infarction models [99, 100]. It is important to note that mouse OSM primarily interacts with the Oncostatin M receptor (OSMR), while human OSM can interact with both OSMR and the leukemia inhibitory factor receptor (LIFR) [65, 101]. In fact, coordinated activation of both OSMR and LIFR signaling is necessary to confer cardioprotection after myocardial infraction in mice [102] and only simultaneous knockdown of both OSMR and LIFR was able to prevent OSM-mediated endothelial cell activation in HUVECs [44]. This suggests that OSM may act upstream of LIFR in human endothelial cells and that dysregulation of LIFR/LIFR expression and signaling connected to OSM activation in HAoEC may promote pathogenic endothelial changes.

Functionally, increased permeability is a hallmark of early vascular dysfunction and facilitates immune cell infiltration into the vessel wall [41, 43]. Consistent with this, OSM-treated HAoEC demonstrated increased FITC-dextran permeability and supported enhanced neutrophil transmigration, indicating that OSM not only disrupts barrier integrity but also promotes functional leukocyte passage. At the molecular level, OSM reduced total occludin protein levels and altered VE-cadherin localization without significantly changing total VE-cadherin abundance. While this may initially appear unexpected, immunofluorescence analysis revealed disrupted junctional organization and increased intracellular extensions of VE-cadherin staining, suggesting redistribution rather than degradation. Homophilic interactions of VE-cadherin in adherens junctions are a major determinant of endothelial barrier integrity and even small focal changes in the molecular organization of cellular junctions can have a functional role to promote gap formation which can result in leakage [103]. Changes in VE-cadherin distribution from a continuous band along cell borders to a serrated zig-zag pattern with extension processes where gaps form in response to different inflammatory stimuli has been reported [104, 105]. These subtle morphological changes representing initial changes in junctional integrity are in line with our observations. Together, these findings indicate that OSM perturbs both tight junction and adherens junction architecture, providing a mechanistic basis for increased permeability and transmigration. OSM has been reported to modulate expression of cell adhesion molecules and disrupt barrier function in skin-equivalent and BBB in vitro models [106, 107], and we extend these findings by demonstrating structural alterations consistent with an inflammatory or proatherogenic endothelial phenotype.

As the subgingival microbial community contains numerous *Treponema* species and based on our previous work demonstrating that *T. denticola* can promote OSM production from neutrophils, we examined whether other oral *Treponema* species can also drive OSM release. While *T. maltophilum* and *T. lecithinolyticum* also elevated OSM release from neutrophils *in vitro* or in a murine air pouch model, *T. denticola* induced the most robust response. Oral *Treponema* spp still had a significantly greater effect than the oral keystone pathogen *P. gingivalis* which did not induce OSM release similar to our previous study [27] or transcription in neutrophils (data not shown), highlighting unique properties of these organisms. A recent report demonstrated that *P. gingivali*s can induce acute kidney injury through OSM/OSMR signaling which suggests tissue and organ context-specific interactions are important to consider [108]. Notably, our findings indicate that OSM secretion from PMNs appears independent of *T. denticola* known virulence factors; such as the major outer sheath membrane protein complex Msp or protease dentilisin. We previously demonstrated that intact spirochetes and secreted products, including purified OMVs, can promote OSM secretion from PMNs [27]. *T. denticola* OMV cargo has been recently characterized [109, 110], and work is ongoing to characterize potential bacterial products and host signaling interactions associated with OSM production. OSM expression and secretion has been reported by microbes in multiple contexts such as human dendritic cells exposed to *E. coli* LPS [111], macrophages or gastric adenocarcinoma cells exposed to *Helicobacter pylori* [112, 113], in mouse lungs during LPS, *Klebsiella* or *E. co*li-induced pneumonia [114, 115] and during human papilloma or cytomegalovirus infection [116, 117]. While LPS can induce expression of OSM in some contexts, this may be organism specific as *Helicobater pylori* induction of OSM appears independent of LPS or the bacterial toxins CagA and VacA [113]. While *T. denticola* lacks an identifiable LPS synthesis pathway [118], a unique lipooligosaccharide (LOS) structure has been reported in the outer membrane of oral *Treponema* species including *T. denticola* [119] and *T. maltophilum* [120] which induce inflammatory response [121–124] which may represent a putative component involved in OSM production from neutrophils.

A strength of this study is the use of a co-culture model to demonstrate that neutrophil-derived OSM is functionally contributing to endothelial cell changes. Conditioned media from *T. denticola*-stimulated neutrophils increased hallmarks of aortic endothelial cell dysfunction including increased permeability, elevated IL-6 production and reduced eNOS levels. Importantly, neutralization of OSM abrogated these effects, supporting a casual role in this model system. Furthermore, preliminary analysis indicates a positive correlation with OSM concentration in conditioned media, supporting a dose-dependent relationship which may corroborate clinical scenarios. Nitric oxide (NO) availability plays an important role in maintaining protective homeostasis in the vasculature by regulating vascular tone, permeability, leukocyte adhesion and platelet aggregation. In endothelial cells, NO is produced primarily by eNOS [125] and reduced eNOS secretion observed in our study represents a hallmark of endothelial dysfunction characterized by dysregulation of NO signaling and can be mechanistically linked to cardiovascular pathologies, due to impaired vasodilation and increased vascular oxidative stress [75, 77, 80]. To our knowledge, the effect of OSM on endothelial NO production remains unknown. However, conflicting data have been reported in microglia with one report demonstrating that OSM increases inducible nitric oxide synthase (iNOS) expression, which is produced under neuroinflammatory conditions, together with NO production in both mouse microglia cell line and primary microglia [126] while another study indicates the opposite effect possibly due to lack of OSM/OSMR expression or handling and experimental differences in the primary cells studied [127]. Our data suggest that OSM may regulate NO production in aortic endothelial cells through modification of eNOS levels, and further work is needed to assess specific molecular processes related to NO generation and eNOS uncoupling underlying endothelial cell dysfunction. Thus, neutrophil-derived OSM not only disrupts structural barrier integrity and promotes inflammatory cytokine production but also suppresses protective endothelial signaling pathways. These findings provide mechanistic evidence that OSM is a principal mediator of endothelial dysfunction in the context of *T. denticola*–neutrophil interaction.

This study utilizes human aortic endothelial cells as an in vitro model to represent a physiologic site prone to adverse vascular events such as atherosclerosis [46], as compared to other commonly used vascular cells lines such as HUVECs as the heterogeneity of EC can affect fundamental understanding of disease conditions [128–131]. Work is also ongoing to examine the effects of OSM signaling and *Treponema* interactions in a physiologically relevant *in vitro* vascular model incorporating smooth muscle cells. The airpouch model supports physiological relevance of *Treponema*-induced OSM release and future studies using models of chronic periodontal infection and vascular disease are important to define long-term consequences. In summary, our findings identify OSM as a central mediator of neutrophil-endothelial crosstalk to promote endothelial cell dysfunction. These results provide mechanistic insight into how periodontal organisms may contribute to vascular inflammation and support OSM signaling as a potential therapeutic target for vascular protection.

## Acknowledgments

We thank Stephen Vanyo for his initial work on this project. Microscopy data in this study was acquired at the Optical Imaging and Analysis Facility, School of Dental Medicine, State University of New York at Buffalo. The spinning disk confocal microscope was supported by an instrumentation grant from The Office of the Director, NIH (S10OD025204) to the State University of New York at Buffalo.

## Notes

### Competing Interest Statement

The authors have declared no competing interest.

## References

1. Lira-Junior R, Boström EA, Gustafsson A. Periodontitis is associated to increased systemic inflammation in postmyocardial infarction patients. Open Heart. 2021;8(2). doi: 10.1136/openhrt-2021-001674. PubMed PMID: 34385358; PubMed Central PMCID: PMCPMC8362710.

2. Thorat M, Pradeep AR, Garg G. Correlation of levels of oncostatin M cytokine in crevicular fluid and serum in periodontal disease. Int J Oral Sci. 2010;2(4):198–207. doi: 10.4248/ijos10077. PubMed PMID: 21404969; PubMed Central PMCID: PMCPMC3499000.

3. Gani DK, Lakshmi D, Krishnan R, Emmadi P. Evaluation of C-reactive protein and interleukin-6 in the peripheral blood of patients with chronic periodontitis. J Indian Soc Periodontol. 2009;13(2):69–74. doi: 10.4103/0972-124x.55840. PubMed PMID: 20407653; PubMed Central PMCID: PMCPMC2847127.

4. Cardoso EM, Reis C, Manzanares-Céspedes MC. Chronic periodontitis, inflammatory cytokines, and interrelationship with other chronic diseases. Postgrad Med. 2018;130(1):98–104. Epub 20171108. doi: 10.1080/00325481.2018.1396876. PubMed PMID: 29065749.

5. Loos BG, Craandijk J, Hoek FJ, Wertheim-van Dillen PM, van der Velden U. Elevation of systemic markers related to cardiovascular diseases in the peripheral blood of periodontitis patients. J Periodontol. 2000;71(10):1528–34. doi: 10.1902/jop.2000.71.10.1528. PubMed PMID: 11063384.

6. Eke PI, Dye BA, Wei L, Slade GD, Thornton-Evans GO, Borgnakke WS, et al. Update on prevalence of periodontitis in adults in the United States: NHANES 2009 to 2012. J Periodontol. 2015;86(5):611–22. doi: 10.1902/jop.2015.140520. PubMed PMID: 25688694; PubMed Central PMCID: PMCPMC4460825.

7. Fu H, Li X, Zhang R, Zhu J, Wang X. Global burden of periodontal diseases among the working-age population from 1990-2021: results from the Global Burden of Disease Study 2021. BMC Public Health. 2025;25(1):1316. Epub 20250408. doi: 10.1186/s12889-025-22566-x. PubMed PMID: 40200262; PubMed Central PMCID: PMCPMC11978096.

8. Nascimento GG, Alves-Costa S, Romandini M. Burden of severe periodontitis and edentulism in 2021, with projections up to 2050: The Global Burden of Disease 2021 study. J Periodontal Res. 2024;59(5):823–67. Epub 20240827. doi: 10.1111/jre.13337. PubMed PMID: 39192495.

9. Page RC, Kornman KS. The pathogenesis of human periodontitis: an introduction. Periodontol 2000. 1997;14:9–11. doi: 10.1111/j.1600-0757.1997.tb00189.x. PubMed PMID: 9567963.

10. Champagne CM, Buchanan W, Reddy MS, Preisser JS, Beck JD, Offenbacher S. Potential for gingival crevice fluid measures as predictors of risk for periodontal diseases. Periodontol 2000. 2003;31:167–80. doi: 10.1034/j.1600-0757.2003.03110.x. PubMed PMID: 12657001.

11. Dewhirst FE, Tamer MA, Ericson RE, Lau CN, Levanos VA, Boches SK, et al. The diversity of periodontal spirochetes by 16S rRNA analysis. Oral Microbiol Immunol. 2000;15(3):196–202. doi: 10.1034/j.1399-302x.2000.150308.x. PubMed PMID: 11154403.

12. Darveau RP. Periodontitis: a polymicrobial disruption of host homeostasis. Nat Rev Microbiol. 2010;8(7):481–90. doi: 10.1038/nrmicro2337. PubMed PMID: 20514045.

13. Choi BK, Paster BJ, Dewhirst FE, Göbel UB. Diversity of cultivable and uncultivable oral spirochetes from a patient with severe destructive periodontitis. Infect Immun. 1994;62(5):1889–95. doi: 10.1128/iai.62.5.1889-1895.1994. PubMed PMID: 8168954; PubMed Central PMCID: PMCPMC186432.

14. Huang D, Wang YY, Li BH, Wu L, Xie WZ, Zhou X, et al. Association between periodontal disease and systemic diseases: a cross-sectional analysis of current evidence. Mil Med Res. 2024;11(1):74. Epub 20241204. doi: 10.1186/s40779-024-00583-y. PubMed PMID: 39633497; PubMed Central PMCID: PMCPMC11616297.

15. Villoria GEM, Fischer RG, Tinoco EMB, Meyle J, Loos BG. Periodontal disease: A systemic condition. Periodontol 2000. 2024;96(1):7–19. Epub 20241104. doi: 10.1111/prd.12616. PubMed PMID: 39494478; PubMed Central PMCID: PMCPMC11579822.

16. Zardawi F, Gul S, Abdulkareem A, Sha A, Yates J. Association Between Periodontal Disease and Atherosclerotic Cardiovascular Diseases: Revisited. Front Cardiovasc Med. 2020;7:625579. Epub 20210115. doi: 10.3389/fcvm.2020.625579. PubMed PMID: 33521070; PubMed Central PMCID: PMCPMC7843501.

17. Larvin H, Kang J, Aggarwal VR, Pavitt S, Wu J. Risk of incident cardiovascular disease in people with periodontal disease: A systematic review and meta-analysis. Clin Exp Dent Res. 2021;7(1):109–22. Epub 20201030. doi: 10.1002/cre2.336. PubMed PMID: 33124761; PubMed Central PMCID: PMCPMC7853902.

18. Chen Y, Rao R, Wu X, Qin Z, Chen Y, Li Q, et al. Periodontitis and the Risk of Heart Failure:a Meta-analysis and Mendelian Randomisation Study. Oral Health Prev Dent. 2025;23:149–64. Epub 20250306. doi: 10.3290/j.ohpd.c_1793. PubMed PMID: 40047704; PubMed Central PMCID: PMCPMC11904829.

19. Roger VL, Go AS, Lloyd-Jones DM, Benjamin EJ, Berry JD, Borden WB, et al. Executive summary: heart disease and stroke statistics--2012 update: a report from the American Heart Association. Circulation. 2012;125(1):188–97. doi: 10.1161/CIR.0b013e3182456d46. PubMed PMID: 22215894.

20. Roth GA, Mensah GA, Johnson CO, Addolorato G, Ammirati E, Baddour LM, et al. Global Burden of Cardiovascular Diseases and Risk Factors, 1990-2019: Update From the GBD 2019 Study. J Am Coll Cardiol. 2020;76(25):2982–3021. doi: 10.1016/j.jacc.2020.11.010. PubMed PMID: 33309175; PubMed Central PMCID: PMCPMC7755038.

21. Darvish M, Shakoor A, Feyz L, Schaap J, van Mieghem NM, de Boer RA, et al. Heart failure: assessment of the global economic burden. Eur Heart J. 2025;46(31):3069–78. doi: 10.1093/eurheartj/ehaf323. PubMed PMID: 40444781; PubMed Central PMCID: PMCPMC12349942.

22. Global, Regional, and National Burden of Cardiovascular Diseases and Risk Factors in 204 Countries and Territories, 1990-2023. J Am Coll Cardiol. 2025;86(22):2167–243. Epub 20250924. doi: 10.1016/j.jacc.2025.08.015. PubMed PMID: 40990886.

23. Papapanou PN. Systemic effects of periodontitis: lessons learned from research on atherosclerotic vascular disease and adverse pregnancy outcomes. Int Dent J. 2015;65(6):283–91. Epub 20150920. doi: 10.1111/idj.12185. PubMed PMID: 26388299; PubMed Central PMCID: PMCPMC4713295.

24. Escobar Arregocés FM, Del Hierro Rada M, Sáenz Martinez MJ, Hernández Meza FJ, Roa NS, Velosa-Porras J, et al. Systemic inflammatory response to non-surgical treatment in hypertensive patients with periodontal infection. Medicine (Baltimore). 2021;100(13):e24951. doi: 10.1097/md.0000000000024951. PubMed PMID: 33787581; PubMed Central PMCID: PMCPMC8021383.

25. Molina A, Ambrosio N, Molina M, Montero E, Virto L, Herrera D, et al. Effect of periodontal therapy on endothelial function and serum biomarkers in patients with periodontitis and established cardiovascular disease: a pilot study. Front Oral Health. 2025;6:1488941. Epub 20250210. doi: 10.3389/froh.2025.1488941. PubMed PMID: 39996093; PubMed Central PMCID: PMCPMC11847872.

26. Tripathy S. The Dual Nature of Oncostatin M: Context-Dependent Mediator of Immunity and Fibrosis. Cell Biochem Biophys. 2025. Epub 20251122. doi: 10.1007/s12013-025-01954-5. PubMed PMID: 41273531.

27. Jones MM, Vanyo ST, Ibraheem W, Maddi A, Visser MB. Treponema denticola stimulates Oncostatin M cytokine release and de novo synthesis in neutrophils and macrophages. J Leukoc Biol. 2020;108(5):1527–41. Epub 2020/07/18. doi: 10.1002/JLB.4MA0620-072RR. PubMed PMID: 32678942.

28. Sonkusle S, Singh V. Comparison of oncostatin M cytokine levels in saliva and serum in periodontitis: a clinicobiochemical study. Can J Dent Hyg. 2024;58(3):155–60. Epub 20241001. PubMed PMID: 39513097; PubMed Central PMCID: PMCPMC11539946.

29. Pradeep AR, S TM, Garima G, Raju A. Serum levels of oncostatin M (a gp 130 cytokine): an inflammatory biomarker in periodontal disease. Biomarkers. 2010;15(3):277–82. doi: 10.3109/13547500903573209. PubMed PMID: 20408777.

30. Ando Y, Tsukasaki M, Huynh NC, Zang S, Yan M, Muro R, et al. The neutrophil-osteogenic cell axis promotes bone destruction in periodontitis. Int J Oral Sci. 2024;16(1):18. Epub 20240227. doi: 10.1038/s41368-023-00275-8. PubMed PMID: 38413562; PubMed Central PMCID: PMCPMC10899642.

31. Matsuoka M, Soria SA, Pires JR, Sant’Ana ACP, Freire M. Natural and induced immune responses in oral cavity and saliva. BMC Immunol. 2025;26(1):34. Epub 20250418. doi: 10.1186/s12865-025-00713-8. PubMed PMID: 40251519; PubMed Central PMCID: PMCPMC12007159.

32. Pothoven KL, Norton JE, Suh LA, Carter RG, Harris KE, Biyasheva A, et al. Neutrophils are a major source of the epithelial barrier disrupting cytokine oncostatin M in patients with mucosal airways disease. J Allergy Clin Immunol. 2017;139(6):1966–78.e9. Epub 20161218. doi: 10.1016/j.jaci.2016.10.039. PubMed PMID: 27993536; PubMed Central PMCID: PMCPMC5529124.

33. Grenier A, Dehoux M, Boutten A, Arce-Vicioso M, Durand G, Gougerot-Pocidalo MA, et al. Oncostatin M production and regulation by human polymorphonuclear neutrophils. Blood. 1999;93(4):1413–21. PubMed PMID: 9949186.

34. Anselmi NK, Vanyo ST, Visser MB. Emerging oral Treponema membrane proteins disorder neutrophil phosphoinositide signaling via phosphatidylinositol-4-phosphate 5-kinase. Front Oral Health. 2025;6:1568983. Epub 20250403. doi: 10.3389/froh.2025.1568983. PubMed PMID: 40248422; PubMed Central PMCID: PMCPMC12003349.

35. Gao L, Kang M, Zhang MJ, Reza Sailani M, Kuraji R, Martinez A, et al. Polymicrobial periodontal disease triggers a wide radius of effect and unique virome. NPJ Biofilms Microbiomes. 2020;6(1):10. Epub 20200310. doi: 10.1038/s41522-020-0120-7. PubMed PMID: 32157085; PubMed Central PMCID: PMCPMC7064479.

36. Askin L, Tanriverdi O, Baris, VO. Oncostatin M and Cardiovascular Diseases: A Narrative Review. Interventional Cardiology Perspectives. 2025;1(1). doi: 10.4274/intercardiopers.2025.2025-2-3.

37. Lindkvist M, Zegeye MM, Grenegård M, Ljungberg LU. Pleiotropic, Unique and Shared Responses Elicited by IL-6 Family Cytokines in Human Vascular Endothelial Cells. Int J Mol Sci. 2022;23(3). Epub 20220127. doi: 10.3390/ijms23031448. PubMed PMID: 35163371; PubMed Central PMCID: PMCPMC8836206.

38. Albasanz-Puig A, Murray J, Preusch M, Coan D, Namekata M, Patel Y, et al. Oncostatin M is expressed in atherosclerotic lesions: a role for Oncostatin M in the pathogenesis of atherosclerosis. Atherosclerosis. 2011;216(2):292–8. Epub 20110303. doi: 10.1016/j.atherosclerosis.2011.02.003. PubMed PMID: 21376322.

39. Jengelley DHA, Wang M, Narasimhan A, Rupert JE, Young AR, Zhong X, et al. Exogenous Oncostatin M induces Cardiac Dysfunction, Musculoskeletal Atrophy, and Fibrosis. Cytokine. 2022;159:155972. Epub 20220830. doi: 10.1016/j.cyto.2022.155972. PubMed PMID: 36054964; PubMed Central PMCID: PMCPMC10468097.

40. Liu C, Wu J, Jia H, Lu C, Liu J, Li Y, et al. Oncostatin M promotes the ox-LDL-induced activation of NLRP3 inflammasomes via the NF-êB pathway in THP-1 macrophages and promotes the progression of atherosclerosis. Ann Transl Med. 2022;10(8):456. doi: 10.21037/atm-22-560. PubMed PMID: 35571419; PubMed Central PMCID: PMCPMC9096425.

41. Naderi-Meshkin H, Setyaningsih WAW. Endothelial Cell Dysfunction: Onset, Progression, and Consequences. Front Biosci (Landmark Ed). 2024;29(6):223. doi: 10.31083/j.fbl2906223. PubMed PMID: 38940049.

42. Chistiakov DA, Orekhov AN, Bobryshev YV. Endothelial Barrier and Its Abnormalities in Cardiovascular Disease. Front Physiol. 2015;6:365. Epub 20151209. doi: 10.3389/fphys.2015.00365. PubMed PMID: 26696899; PubMed Central PMCID: PMCPMC4673665.

43. Mudau M, Genis A, Lochner A, Strijdom H. Endothelial dysfunction: the early predictor of atherosclerosis. Cardiovasc J Afr. 2012;23(4):222–31. doi: 10.5830/cvja-2011-068. PubMed PMID: 22614668; PubMed Central PMCID: PMCPMC3721957.

44. van Keulen D, Pouwer MG, Pasterkamp G, van Gool AJ, Sollewijn Gelpke MD, Princen HMG, et al. Inflammatory cytokine oncostatin M induces endothelial activation in macro- and microvascular endothelial cells and in APOE*3Leiden.CETP mice. PLoS One. 2018;13(10):e0204911. Epub 20181001. doi: 10.1371/journal.pone.0204911. PubMed PMID: 30273401; PubMed Central PMCID: PMCPMC6166945.

45. Setiadi H, Yago T, Liu Z, McEver RP. Endothelial signaling by neutrophil-released oncostatin M enhances P-selectin-dependent inflammation and thrombosis. Blood Adv. 2019;3(2):168–83. doi: 10.1182/bloodadvances.2018026294. PubMed PMID: 30670533; PubMed Central PMCID: PMCPMC6341191.

46. Aboyans V, Lacroix P, Criqui MH. Large and small vessels atherosclerosis: similarities and differences. Prog Cardiovasc Dis. 2007;50(2):112–25. doi: 10.1016/j.pcad.2007.04.001. PubMed PMID: 17765473.

47. Kitajima S, Sakuma S, Morimoto M. Macroscopic distribution of coronary atherosclerotic lesions in cholesterol-fed rabbits. Exp Anim. 1998;47(4):221–7. doi: 10.1538/expanim.47.221. PubMed PMID: 10067164.

48. Falk E. Pathogenesis of atherosclerosis. J Am Coll Cardiol. 2006;47(8 Suppl):C7–12. doi: 10.1016/j.jacc.2005.09.068. PubMed PMID: 16631513.

49. Fenno JC. Laboratory maintenance of Treponema denticola. Curr Protoc Microbiol. 2005;Chapter 12:Unit 12B.1. doi: 10.1002/9780471729259.mc12b01s00. PubMed PMID: 18770551.

50. Anselmi NK, Bynum K, Kay JG, Visser MB. Analysis of Neutrophil Responses to Biological Exposures. Curr Protoc. 2023;3(6):e827. doi: 10.1002/cpz1.827. PubMed PMID: 37358215; PubMed Central PMCID: PMCPMC10416710.

51. Lakschevitz FS, Glogauer M. High-purity neutrophil isolation from human peripheral blood and saliva for transcriptome analysis. Methods Mol Biol. 2014;1124:469–83. doi: 10.1007/978-1-62703-845-4_28. PubMed PMID: 24504969.

52. He X, Jiang W, Luo Z, Qu T, Wang Z, Liu N, et al. IFN-ã regulates human dental pulp stem cells behavior via NF-êB and MAPK signaling. Sci Rep. 2017;7:40681. Epub 20170118. doi: 10.1038/srep40681. PubMed PMID: 28098169; PubMed Central PMCID: PMCPMC5241669.

53. Ito F, Mori T, Takaoka O, Tanaka Y, Koshiba A, Tatsumi H, et al. Effects of drospirenone on adhesion molecule expression and monocyte adherence in human endothelial cells. Eur J Obstet Gynecol Reprod Biol. 2016;201:113–7. Epub 20160407. doi: 10.1016/j.ejogrb.2016.03.044. PubMed PMID: 27088625.

54. Bi J, Intriago MFB, Koivisto L, Jiang G, Häkkinen L, Larjava H. Leucocyte- and platelet-rich fibrin regulates expression of genes related to early wound healing in human gingival fibroblasts. J Clin Periodontol. 2020;47(7):851–62. Epub 20200513. doi: 10.1111/jcpe.13293. PubMed PMID: 32304115.

55. Lakschevitz FS, Aboodi GM, Glogauer M. Oral neutrophils display a site-specific phenotype characterized by expression of T-cell receptors. J Periodontol. 2013;84(10):1493–503. doi: 10.1902/jop.2012.120477. PubMed PMID: 23205919.

56. Bigildeev AE, Zezina EA, Shipounova IN, Drize NJ. Interleukin-1 beta enhances human multipotent mesenchymal stromal cell proliferative potential and their ability to maintain hematopoietic precursor cells. Cytokine. 2015;71(2):246–54. Epub 20141124. doi: 10.1016/j.cyto.2014.10.018. PubMed PMID: 25461405.

57. Anselmi NK, Vanyo ST, Clark ND, Rodriguez DML, Jones MM, Rosenthal S, et al. Topology and functional characterization of major outer membrane proteins of Treponema maltophilum and Treponema lecithinolyticum. Mol Oral Microbiol. 2025;40(1):17–36. Epub 20240912. doi: 10.1111/omi.12484. PubMed PMID: 39263909; PubMed Central PMCID: PMCPMC11752107.

58. Gruson D, Ferracin B, Ahn SA, Rousseau MF. Elevation of plasma oncostatin M in heart failure. Future Cardiol. 2017;13(3):219–27. Epub 20170606. doi: 10.2217/fca-2016-0063. PubMed PMID: 28585906.

59. Ikeda S, Sato K, Takeda M, Miki K, Aizawa K, Takada T, et al. Oncostatin M is a novel biomarker for coronary artery disease - A possibility as a screening tool of silent myocardial ischemia for diabetes mellitus. Int J Cardiol Heart Vasc. 2021;35:100829. Epub 20210626. doi: 10.1016/j.ijcha.2021.100829. PubMed PMID: 34235245; PubMed Central PMCID: PMCPMC8250159.

60. Neele AE, Chen HJ, Gijbels MJJ, van der Velden S, Hoeksema MA, Boshuizen MCS, et al. Myeloid Ezh2 Deficiency Limits Atherosclerosis Development. Front Immunol. 2020;11:594603. Epub 20210126. doi: 10.3389/fimmu.2020.594603. PubMed PMID: 33574814; PubMed Central PMCID: PMCPMC7871783.

61. Rohde D, Vandoorne K, Lee IH, Grune J, Zhang S, McAlpine CS, et al. Bone marrow endothelial dysfunction promotes myeloid cell expansion in cardiovascular disease. Nat Cardiovasc Res. 2022;1(1):28–44. Epub 20211223. doi: 10.1038/s44161-021-00002-8. PubMed PMID: 35747128; PubMed Central PMCID: PMCPMC9216333.

62. Sumagin R, Lomakina E, Sarelius IH. Leukocyte-endothelial cell interactions are linked to vascular permeability via ICAM-1-mediated signaling. Am J Physiol Heart Circ Physiol. 2008;295(3):H969–h77. Epub 20080718. doi: 10.1152/ajpheart.00400.2008. PubMed PMID: 18641276; PubMed Central PMCID: PMCPMC2544502.

63. Liu G, Place AT, Chen Z, Brovkovych VM, Vogel SM, Muller WA, et al. ICAM-1-activated Src and eNOS signaling increase endothelial cell surface PECAM-1 adhesivity and neutrophil transmigration. Blood. 2012;120(9):1942–52. Epub 20120717. doi: 10.1182/blood-2011-12-397430. PubMed PMID: 22806890; PubMed Central PMCID: PMCPMC3433096.

64. Marzolla V, Armani A, Mammi C, Moss ME, Pagliarini V, Pontecorvo L, et al. Essential role of ICAM-1 in aldosterone-induced atherosclerosis. Int J Cardiol. 2017;232:233–42. Epub 20170105. doi: 10.1016/j.ijcard.2017.01.013. PubMed PMID: 28089144; PubMed Central PMCID: PMCPMC5890338.

65. Zhou Y, Stevis PE, Cao J, Ehrlich G, Jones J, Rafique A, et al. Structures of complete extracellular assemblies of type I and type II Oncostatin M receptor complexes. Nat Commun. 2024;15(1):9776. Epub 20241112. doi: 10.1038/s41467-024-54124-1. PubMed PMID: 39532904; PubMed Central PMCID: PMCPMC11557873.

66. Hemme E, Depuydt MAC, van Santbrink PJ, Wezel A, Smeets HJ, Foks AC, et al. Leukemia inhibitory factor receptor inhibition by EC359 reduces atherosclerotic stenosis grade in Ldlr(-/-) mice. Eur J Pharmacol. 2024;985:177121. Epub 20241109. doi: 10.1016/j.ejphar.2024.177121. PubMed PMID: 39528103.

67. Zhang X, Li J, Qin JJ, Cheng WL, Zhu X, Gong FH, et al. Oncostatin M receptor β deficiency attenuates atherogenesis by inhibiting JAK2/STAT3 signaling in macrophages. J Lipid Res. 2017;58(5):895–906. Epub 20170303. doi: 10.1194/jlr.M074112. PubMed PMID: 28258089; PubMed Central PMCID: PMCPMC5408608.

68. Claesson-Welsh L, Dejana E, McDonald DM. Permeability of the Endothelial Barrier: Identifying and Reconciling Controversies. Trends Mol Med. 2021;27(4):314–31. Epub 20201210. doi: 10.1016/j.molmed.2020.11.006. PubMed PMID: 33309601; PubMed Central PMCID: PMCPMC8005435.

69. Allport JR, Ding H, Collins T, Gerritsen ME, Luscinskas FW. Endothelial-dependent mechanisms regulate leukocyte transmigration: a process involving the proteasome and disruption of the vascular endothelial-cadherin complex at endothelial cell-to-cell junctions. J Exp Med. 1997;186(4):517–27. doi: 10.1084/jem.186.4.517. PubMed PMID: 9254650; PubMed Central PMCID: PMCPMC2199034.

70. Angelini DJ, Hyun SW, Grigoryev DN, Garg P, Gong P, Singh IS, et al. TNF-alpha increases tyrosine phosphorylation of vascular endothelial cadherin and opens the paracellular pathway through fyn activation in human lung endothelia. Am J Physiol Lung Cell Mol Physiol. 2006;291(6):L1232–45. Epub 20060804. doi: 10.1152/ajplung.00109.2006. PubMed PMID: 16891393.

71. Griffen AL, Beall CJ, Campbell JH, Firestone ND, Kumar PS, Yang ZK, et al. Distinct and complex bacterial profiles in human periodontitis and health revealed by 16S pyrosequencing. Isme j. 2012;6(6):1176–85. Epub 20111215. doi: 10.1038/ismej.2011.191. PubMed PMID: 22170420; PubMed Central PMCID: PMCPMC3358035.

72. Visser MB, Ellen RP. New insights into the emerging role of oral spirochaetes in periodontal disease. Clin Microbiol Infect. 2011;17(4):502–12. Epub 2011/03/19. doi: 10.1111/j.1469-0691.2011.03460.x. PubMed PMID: 21414084.

73. Ghosh S, Tuz AA, Stenzel M, Singh V, Richter M, Soehnlein O, et al. Proteomic Characterization of 1000 Human and Murine Neutrophils Freshly Isolated From Blood and Sites of Sterile Inflammation. Mol Cell Proteomics. 2024;23(11):100858. Epub 20241011. doi: 10.1016/j.mcpro.2024.100858. PubMed PMID: 39395581; PubMed Central PMCID: PMCPMC11630641.

74. Sollberger G, Brenes AJ, Warner J, Arthur JSC, Howden AJM. Quantitative proteomics reveals tissue-specific, infection-induced and species-specific neutrophil protein signatures. Sci Rep. 2024;14(1):5966. Epub 20240312. doi: 10.1038/s41598-024-56163-6. PubMed PMID: 38472281; PubMed Central PMCID: PMCPMC10933280.

75. Tran N, Garcia T, Aniqa M, Ali S, Ally A, Nauli SM. Endothelial Nitric Oxide Synthase (eNOS) and the Cardiovascular System: in Physiology and in Disease States. Am J Biomed Sci Res. 2022;15(2):153–77. Epub 20220104. PubMed PMID: 35072089; PubMed Central PMCID: PMCPMC8774925.

76. Janaszak-Jasiecka A, Płoska A, Wieroñska JM, Dobrucki LW, Kalinowski L. Endothelial dysfunction due to eNOS uncoupling: molecular mechanisms as potential therapeutic targets. Cell Mol Biol Lett. 2023;28(1):21. Epub 20230309. doi: 10.1186/s11658-023-00423-2. PubMed PMID: 36890458; PubMed Central PMCID: PMCPMC9996905.

77. Kawashima S, Yokoyama M. Dysfunction of endothelial nitric oxide synthase and atherosclerosis. Arterioscler Thromb Vasc Biol. 2004;24(6):998–1005. Epub 20040304. doi: 10.1161/01.Atv.0000125114.88079.96. PubMed PMID: 15001455.

78. Smith AR, Visioli F, Hagen TM. Plasma membrane-associated endothelial nitric oxide synthase and activity in aging rat aortic vascular endothelia markedly decline with age. Arch Biochem Biophys. 2006;454(1):100–5. Epub 20060309. doi: 10.1016/j.abb.2006.02.017. PubMed PMID: 16982030.

79. Barberà JA, Peinado VI, Santos S, Ramirez J, Roca J, Rodriguez-Roisin R. Reduced expression of endothelial nitric oxide synthase in pulmonary arteries of smokers. Am J Respir Crit Care Med. 2001;164(4):709–13. doi: 10.1164/ajrccm.164.4.2101023. PubMed PMID: 11520741.

80. Oemar BS, Tschudi MR, Godoy N, Brovkovich V, Malinski T, Lüscher TF. Reduced endothelial nitric oxide synthase expression and production in human atherosclerosis. Circulation. 1998;97(25):2494–8. doi: 10.1161/01.cir.97.25.2494. PubMed PMID: 9657467.

81. Modur V, Feldhaus MJ, Weyrich AS, Jicha DL, Prescott SM, Zimmerman GA, et al. Oncostatin M is a proinflammatory mediator. In vivo effects correlate with endothelial cell expression of inflammatory cytokines and adhesion molecules. J Clin Invest. 1997;100(1):158–68. doi: 10.1172/jci119508. PubMed PMID: 9202068; PubMed Central PMCID: PMCPMC508176.

82. Bevilacqua MP, Stengelin S, Gimbrone MA, Jr., Seed B. Endothelial leukocyte adhesion molecule 1: an inducible receptor for neutrophils related to complement regulatory proteins and lectins. Science. 1989;243(4895):1160–5. doi: 10.1126/science.2466335. PubMed PMID: 2466335.

83. Phillips ML, Nudelman E, Gaeta FC, Perez M, Singhal AK, Hakomori S, et al. ELAM-1 mediates cell adhesion by recognition of a carbohydrate ligand, sialyl-Lex. Science. 1990;250(4984):1130–2. doi: 10.1126/science.1701274. PubMed PMID: 1701274.

84. Zhang J, Huang S, Zhu Z, Gatt A, Liu J. E-selectin in vascular pathophysiology. Front Immunol. 2024;15:1401399. Epub 20240719. doi: 10.3389/fimmu.2024.1401399. PubMed PMID: 39100681; PubMed Central PMCID: PMCPMC11294169.

85. Kerfoot SM, Raharjo E, Ho M, Kaur J, Serirom S, McCafferty DM, et al. Exclusive neutrophil recruitment with oncostatin M in a human system. Am J Pathol. 2001;159(4):1531–9. doi: 10.1016/s0002-9440(10)62538-2. PubMed PMID: 11583979; PubMed Central PMCID: PMCPMC1850489.

86. Yang L, Froio RM, Sciuto TE, Dvorak AM, Alon R, Luscinskas FW. ICAM-1 regulates neutrophil adhesion and transcellular migration of TNF-alpha-activated vascular endothelium under flow. Blood. 2005;106(2):584–92. Epub 20050405. doi: 10.1182/blood-2004-12-4942. PubMed PMID: 15811956; PubMed Central PMCID: PMCPMC1635241.

87. van der Meer IM, de Maat MP, Bots ML, Breteler MM, Meijer J, Kiliaan AJ, et al. Inflammatory mediators and cell adhesion molecules as indicators of severity of atherosclerosis: the Rotterdam Study. Arterioscler Thromb Vasc Biol. 2002;22(5):838–42. doi: 10.1161/01.atv.0000016249.96529.b8. PubMed PMID: 12006399.

88. Ridker PM, Hennekens CH, Roitman-Johnson B, Stampfer MJ, Allen J. Plasma concentration of soluble intercellular adhesion molecule 1 and risks of future myocardial infarction in apparently healthy men. Lancet. 1998;351(9096):88–92. doi: 10.1016/s0140-6736(97)09032-6. PubMed PMID: 9439492.

89. Ruprecht K, Kuhlmann T, Seif F, Hummel V, Kruse N, Brück W, et al. Effects of oncostatin M on human cerebral endothelial cells and expression in inflammatory brain lesions. J Neuropathol Exp Neurol. 2001;60(11):1087–98. doi: 10.1093/jnen/60.11.1087. PubMed PMID: 11706938.

90. Schnittker D, Kwofie K, Ashkar A, Trigatti B, Richards CD. Oncostatin M and TLR-4 ligand synergize to induce MCP-1, IL-6, and VEGF in human aortic adventitial fibroblasts and smooth muscle cells. Mediators Inflamm. 2013;2013:317503. Epub 20131106. doi: 10.1155/2013/317503. PubMed PMID: 24307759; PubMed Central PMCID: PMCPMC3836373.

91. Bumm CV, Schwendicke F, Heck K, Frasheri I, Summer B, Ern C, et al. The Role of Interleukin-8 in the Estimation of Responsiveness to Steps 1 and 2 of Periodontal Therapy. J Clin Periodontol. 2024;51(11):1433–42. Epub 20240816. doi: 10.1111/jcpe.14055. PubMed PMID: 39152683.

92. Finoti LS, Nepomuceno R, Pigossi SC, Corbi SC, Secolin R, Scarel-Caminaga RM. Association between interleukin-8 levels and chronic periodontal disease: A PRISMA-compliant systematic review and meta-analysis. Medicine (Baltimore). 2017;96(22):e6932. doi: 10.1097/md.0000000000006932. PubMed PMID: 28562542; PubMed Central PMCID: PMCPMC5459707.

93. Jin L, Söder B, Corbet EF. Interleukin-8 and granulocyte elastase in gingival crevicular fluid in relation to periodontopathogens in untreated adult periodontitis. J Periodontol. 2000;71(6):929–39. doi: 10.1902/jop.2000.71.6.929. PubMed PMID: 10914796.

94. Hiyoshi T, Domon H, Maekawa T, Tamura H, Isono T, Hirayama S, et al. Neutrophil elastase aggravates periodontitis by disrupting gingival epithelial barrier via cleaving cell adhesion molecules. Sci Rep. 2022;12(1):8159. Epub 20220517. doi: 10.1038/s41598-022-12358-3. PubMed PMID: 35581391; PubMed Central PMCID: PMCPMC9114116.

95. Ahmad P, Slots J, Siqueira WL. Serum cytokines in periodontal diseases. Periodontol 2000. 2025;98(1):138–80. Epub 20250925. doi: 10.1111/prd.12629. PubMed PMID: 40995683; PubMed Central PMCID: PMCPMC12842884.

96. Zhang X, Chen S, Yin G, Liang P, Feng Y, Yu W, et al. The Role of JAK/STAT Signaling Pathway and Its Downstream Influencing Factors in the Treatment of Atherosclerosis. J Cardiovasc Pharmacol Ther. 2024;29:10742484241248046. doi: 10.1177/10742484241248046. PubMed PMID: 38656132.

97. Wang R, Zhang Y, Xu L, Lin Y, Yang X, Bai L, et al. Protein Inhibitor of Activated STAT3 Suppresses Oxidized LDL-induced Cell Responses during Atherosclerosis in Apolipoprotein E-deficient Mice. Sci Rep. 2016;6:36790. Epub 20161115. doi: 10.1038/srep36790. PubMed PMID: 27845432; PubMed Central PMCID: PMCPMC5109228.

98. Chen Q, Lv J, Yang W, Xu B, Wang Z, Yu Z, et al. Targeted inhibition of STAT3 as a potential treatment strategy for atherosclerosis. Theranostics. 2019;9(22):6424–42. Epub 20190814. doi: 10.7150/thno.35528. PubMed PMID: 31588227; PubMed Central PMCID: PMCPMC6771242.

99. Berry MF, Pirolli TJ, Jayasankar V, Morine KJ, Moise MA, Fisher O, et al. Targeted overexpression of leukemia inhibitory factor to preserve myocardium in a rat model of postinfarction heart failure. J Thorac Cardiovasc Surg. 2004;128(6):866–75. doi: 10.1016/j.jtcvs.2004.06.046. PubMed PMID: 15573071.

100. Kanda M, Nagai T, Takahashi T, Liu ML, Kondou N, Naito AT, et al. Leukemia Inhibitory Factor Enhances Endogenous Cardiomyocyte Regeneration after Myocardial Infarction. PLoS One. 2016;11(5):e0156562. Epub 20160526. doi: 10.1371/journal.pone.0156562. PubMed PMID: 27227407; PubMed Central PMCID: PMCPMC4881916.

101. Adrian-Segarra JM, Sreenivasan K, Gajawada P, Lörchner H, Braun T, Pöling J. The AB loop of oncostatin M (OSM) determines species-specific signaling in humans and mice. J Biol Chem. 2018;293(52):20181–99. Epub 20181029. doi: 10.1074/jbc.RA118.004375. PubMed PMID: 30373773; PubMed Central PMCID: PMCPMC6311520.

102. Lörchner H, Adrian-Segarra JM, Waechter C, Wagner R, Góes ME, Brachmann N, et al. Concomitant Activation of OSM and LIF Receptor by a Dual-Specific hlOSM Variant Confers Cardioprotection after Myocardial Infarction in Mice. Int J Mol Sci. 2021;23(1). Epub 20211229. doi: 10.3390/ijms23010353. PubMed PMID: 35008777; PubMed Central PMCID: PMCPMC8745562.

103. Harris ES, Nelson WJ. VE-cadherin: at the front, center, and sides of endothelial cell organization and function. Curr Opin Cell Biol. 2010;22(5):651–8. Epub 20100811. doi: 10.1016/j.ceb.2010.07.006. PubMed PMID: 20708398; PubMed Central PMCID: PMCPMC2948582.

104. Postma RJ, Fischer SE, Bijkerk R, van Zonneveld AJ. Unveiling endothelial cell border heterogeneity: VE-cadherin adherens junction stratification by deep convolutional neural networks. PLoS One. 2025;20(1):e0317110. Epub 20250106. doi: 10.1371/journal.pone.0317110. PubMed PMID: 39761260; PubMed Central PMCID: PMCPMC11703098.

105. Li X, Padhan N, Sjöström EO, Roche FP, Testini C, Honkura N, et al. VEGFR2 pY949 signalling regulates adherens junction integrity and metastatic spread. Nat Commun. 2016;7:11017. Epub 20160323. doi: 10.1038/ncomms11017. PubMed PMID: 27005951; PubMed Central PMCID: PMCPMC4814575.

106. Hermans D, Houben E, Baeten P, Slaets H, Janssens K, Hoeks C, et al. Oncostatin M triggers brain inflammation by compromising blood-brain barrier integrity. Acta Neuropathol. 2022;144(2):259–81. Epub 20220606. doi: 10.1007/s00401-022-02445-0. PubMed PMID: 35666306.

107. Gazel A, Rosdy M, Bertino B, Tornier C, Sahuc F, Blumenberg M. A characteristic subset of psoriasis-associated genes is induced by oncostatin-M in reconstituted epidermis. J Invest Dermatol. 2006;126(12):2647–57. Epub 20060817. doi: 10.1038/sj.jid.5700461. PubMed PMID: 16917497.

108. Wei W, Sun J, Ji Z, Hu J, Jiang Q. Gingipain and oncostatin M synergistically disrupt kidney tight junctions in periodontitis-associated acute kidney injury. J Periodontol. 2024;95(9):867–79. Epub 20240704. doi: 10.1002/jper.24-0007. PubMed PMID: 38963713.

109. Veith PD, Glew MD, Gorasia DG, Chen D, O’Brien-Simpson NM, Reynolds EC. Localization of Outer Membrane Proteins in Treponema denticola by Quantitative Proteome Analyses of Outer Membrane Vesicles and Cellular Fractions. J Proteome Res. 2019;18(4):1567–81. Epub 20190226. doi: 10.1021/acs.jproteome.8b00860. PubMed PMID: 30761904.

110. Jones MM, Vanyo ST, Visser MB. The Msp protein of Treponema denticola interrupts activity of phosphoinositide processing in neutrophils. Infect Immun. 2019. Epub 2019/09/05. doi: 10.1128/IAI.00553-19. PubMed PMID: 31481407.

111. Suda T, Chida K, Todate A, Ide K, Asada K, Nakamura Y, et al. Oncostatin M production by human dendritic cells in response to bacterial products. Cytokine. 2002;17(6):335–40. doi: 10.1006/cyto.2002.1023. PubMed PMID: 12061841.

112. Zeaiter Z, Diaz H, Stein M, Huynh HQ. Helicobacter pylori induces expression and secretion of oncostatin M in macrophages in vitro. Dig Dis Sci. 2011;56(3):689–97. Epub 20100727. doi: 10.1007/s10620-010-1341-z. PubMed PMID: 20661773.

113. Zoaiter M, Nasser R, Hage-Sleiman R, Abdel-Sater F, Badran B, Zeaiter Z. Helicobacter pylori outer membrane vesicles induce expression and secretion of oncostatin M in AGS gastric cancer cells. Braz J Microbiol. 2021;52(3):1057–66. Epub 20210413. doi: 10.1007/s42770-021-00490-7. PubMed PMID: 33851342; PubMed Central PMCID: PMCPMC8324738.

114. Headland SE, Dengler HS, Xu D, Teng G, Everett C, Ratsimandresy RA, et al. Oncostatin M expression induced by bacterial triggers drives airway inflammatory and mucus secretion in severe asthma. Sci Transl Med. 2022;14(627):eabf8188. Epub 20220112. doi: 10.1126/scitranslmed.abf8188. PubMed PMID: 35020406.

115. Quinton LJ, Jones MR, Robson BE, Simms BT, Whitsett JA, Mizgerd JP. Alveolar epithelial STAT3, IL-6 family cytokines, and host defense during Escherichia coli pneumonia. Am J Respir Cell Mol Biol. 2008;38(6):699–706. Epub 20080110. doi: 10.1165/rcmb.2007-0365OC. PubMed PMID: 18192501; PubMed Central PMCID: PMCPMC2396249.

116. Dumortier J, Streblow DN, Moses AV, Jacobs JM, Kreklywich CN, Camp D, et al. Human cytomegalovirus secretome contains factors that induce angiogenesis and wound healing. J Virol. 2008;82(13):6524–35. Epub 20080430. doi: 10.1128/jvi.00502-08. PubMed PMID: 18448536; PubMed Central PMCID: PMCPMC2447085.

117. Chan G, Bivins-Smith ER, Smith MS, Smith PM, Yurochko AD. Transcriptome analysis reveals human cytomegalovirus reprograms monocyte differentiation toward an M1 macrophage. J Immunol. 2008;181(1):698–711. doi: 10.4049/jimmunol.181.1.698. PubMed PMID: 18566437; PubMed Central PMCID: PMCPMC2614917.

118. Seshadri R, Myers GS, Tettelin H, Eisen JA, Heidelberg JF, Dodson RJ, et al. Comparison of the genome of the oral pathogen Treponema denticola with other spirochete genomes. Proc Natl Acad Sci U S A. 2004;101(15):5646–51. Epub 20040402. doi: 10.1073/pnas.0307639101. PubMed PMID: 15064399; PubMed Central PMCID: PMCPMC397461.

119. Schultz CP, Wolf V, Lange R, Mertens E, Wecke J, Naumann D, et al. Evidence for a new type of outer membrane lipid in oral spirochete Treponema denticola. Functioning permeation barrier without lipopolysaccharides. J Biol Chem. 1998;273(25):15661–6. doi: 10.1074/jbc.273.25.15661. PubMed PMID: 9624160.

120. Opitz B, Schröder NW, Spreitzer I, Michelsen KS, Kirschning CJ, Hallatschek W, et al. Toll-like receptor-2 mediates Treponema glycolipid and lipoteichoic acid-induced NF-kappaB translocation. J Biol Chem. 2001;276(25):22041–7. Epub 20010402. doi: 10.1074/jbc.M010481200. PubMed PMID: 11285258.

121. Rosen G, Sela MN, Naor R, Halabi A, Barak V, Shapira L. Activation of murine macrophages by lipoprotein and lipooligosaccharide of Treponema denticola. Infect Immun. 1999;67(3):1180–6. doi: 10.1128/iai.67.3.1180-1186.1999. PubMed PMID: 10024558; PubMed Central PMCID: PMCPMC96444.

122. Nussbaum G, Ben-Adi S, Genzler T, Sela M, Rosen G. Involvement of Toll-like receptors 2 and 4 in the innate immune response to Treponema denticola and its outer sheath components. Infect Immun. 2009;77(9):3939–47. Epub 20090713. doi: 10.1128/iai.00488-09. PubMed PMID: 19596768; PubMed Central PMCID: PMCPMC2738032.

123. Choi BK, Lee HJ, Kang JH, Jeong GJ, Min CK, Yoo YJ. Induction of osteoclastogenesis and matrix metalloproteinase expression by the lipooligosaccharide of Treponema denticola. Infect Immun. 2003;71(1):226–33. doi: 10.1128/iai.71.1.226-233.2003. PubMed PMID: 12496170; PubMed Central PMCID: PMCPMC143274.

124. Schröder NW, Opitz B, Lamping N, Michelsen KS, Zähringer U, Göbel UB, et al. Involvement of lipopolysaccharide binding protein, CD14, and Toll-like receptors in the initiation of innate immune responses by Treponema glycolipids. J Immunol. 2000;165(5):2683–93. doi: 10.4049/jimmunol.165.5.2683. PubMed PMID: 10946299.

125. Förstermann U, Sessa WC. Nitric oxide synthases: regulation and function. Eur Heart J. 2012;33(7):829–37, 37a-37d. Epub 20110901. doi: 10.1093/eurheartj/ehr304. PubMed PMID: 21890489; PubMed Central PMCID: PMCPMC3345541.

126. Baker BJ, Park KW, Qin H, Ma X, Benveniste EN. IL-27 inhibits OSM-mediated TNF-alpha and iNOS gene expression in microglia. Glia. 2010;58(9):1082–93. doi: 10.1002/glia.20989. PubMed PMID: 20468050; PubMed Central PMCID: PMCPMC3378052.

127. Hsu MP, Frausto R, Rose-John S, Campbell IL. Analysis of IL-6/gp130 family receptor expression reveals that in contrast to astroglia, microglia lack the oncostatin M receptor and functional responses to oncostatin M. Glia. 2015;63(1):132–41. Epub 20140807. doi: 10.1002/glia.22739. PubMed PMID: 25103368.

128. Gomez-Salinero JM, Redmond D, Rafii S. Microenvironmental determinants of endothelial cell heterogeneity. Nat Rev Mol Cell Biol. 2025;26(6):476–95. Epub 20250128. doi: 10.1038/s41580-024-00825-w. PubMed PMID: 39875728.

129. Seo HR, Jeong HE, Joo HJ, Choi SC, Park CY, Kim JH, et al. Intrinsic FGF2 and FGF5 promotes angiogenesis of human aortic endothelial cells in 3D microfluidic angiogenesis system. Sci Rep. 2016;6:28832. Epub 20160630. doi: 10.1038/srep28832. PubMed PMID: 27357248; PubMed Central PMCID: PMCPMC4928073.

130. Morris TE, Mattox PA, Shipley GD, Wagner CR, Hosenpud JD. The pattern of cytokine messenger RNA expression in human aortic endothelial cells is different from that of human umbilical vein endothelial cells. Transpl Immunol. 1993;1(2):137–42. doi: 10.1016/0966-3274(93)90007-u. PubMed PMID: 7915952.

131. Sung HJ, Yee A, Eskin SG, McIntire LV. Cyclic strain and motion control produce opposite oxidative responses in two human endothelial cell types. Am J Physiol Cell Physiol. 2007;293(1):C87–94. Epub 20070221. doi: 10.1152/ajpcell.00585.2006. PubMed PMID: 17314265.

